# Wavelet transform of single-trial vestibular short-latency evoked potential reveals temporary reduction in signal detectability and temporal precision following noise exposure

**DOI:** 10.1101/2024.06.10.596660

**Authors:** Mamiko Niwa, David Bauer, Marie Anderson, Ariane Kanicki, Richard A. Altschuler, Courtney E. Stewart, W. Michael King

## Abstract

The vestibular short-latency evoked potential (VsEP) reflects the activity of irregular vestibular afferents and their target neurons in the brain stem. Attenuation of trial-averaged VsEP waveforms is widely accepted as an indicator of vestibular dysfunction, however, more quantitative analyses of VsEP waveforms could reveal underlying neural properties of VsEP waveforms. Here, we present a time-frequency analysis of the VsEP with a wavelet transform on a single-trial basis, which allows us to examine trial-by-trial variability in the strength of VsEP waves as well as their temporal coherence across trials. Using this method, we examined changes in the VsEP following 110 dB SPL noise exposure in rats. We found detectability of head jerks based on the power of wavelet transform coefficients was significantly reduced 1 day after noise exposure but recovered nearly to pre-exposure level in 3 - 7 days and completely by 28 days after exposure. Temporal coherence of VsEP waves across trials was also significantly reduced on 1 day after exposure but recovered with a similar time course. Additionally, we found a significant reduction in the number of calretinin-positive calyces in the sacculi collected 28 days after noise exposure. Furthermore, the number of calretinin-positive calyces was significantly correlated with the degree of reduction in temporal coherence and/or signal detectability of the smallest-amplitude jerks. This new analysis of the VsEP provides more quantitative descriptions of noise-induced changes as well as new insights into potential mechanisms underlying noise-induced vestibular dysfunction.

**Significance Statement:** Our study presents a new method of VsEP quantification using wavelet transform on a single-trial basis. It also describes a novel approach to determine the stimulus threshold of the VsEP based on signal-detection theory and Rayleigh statistics. The present analysis could also be applied to analysis of auditory brain stem response (ABR). Thus, it has the potential to provide new insights into the physiological properties that underlie peripheral vestibular and auditory dysfunction.

## Introduction

Vestibular short-latency evoked potential (VsEP) recording has been widely used as a non-invasive, physiological test of irregular vestibular afferent activity. The VsEP, when elicited by a ramp of linear acceleration, is thought to represent the evoked activity of irregular afferents innervating the saccule and utricle, vestibular sensory organs that sense linear and gravitational acceleration of an animal’s head. The VsEP has traditionally been analyzed by averaging responses to repeated stimulations and identifying peaks on a trial-averaged VsEP waveform. Reduction in peak amplitude or prolonged peak latency has been reported in animal models of vestibular dysfunction associated with genetic (Ono et al., 2020), iatrogenic (e.g., Bremer et al., 2014), and environmental factors (e.g., Stewart et al., 2020). Thus, the VsEP is used as an indicator of reduced and/or lost vestibular function. Despite the wide use of the VsEP and its capability of capturing gross changes in the vestibular system, its quantification based on the peaks of a trial-averaged waveform is limited.

Here, we describe a time-frequency analysis of the VsEP waveform that uses a wavelet transform on a single-trial basis. The wavelet transform exposes the frequency content of the VsEP waveform and how the frequency spectrum changes over time. By applying the transform on a single-trial basis, we are able to examine the trial-by-trial variability in the strength of the underlying VsEP waves as well as the trial-by-trial coherence in the timing of these waves. This information cannot be gained by examining a trial-averaged VsEP waveform.

Additionally, we provide a statistical approach to determine the stimulus threshold of the VsEP. Determination of a threshold stimulus level based on the shape of the waveform may vary between observers and is difficult to determine if the pattern of the VsEP waveform is atypical due to pathological changes in an animal’s vestibular periphery. In this study, we determine thresholds based on the power and phase of wavelet transform coefficients using the area under the receiver operating characteristics curve (ROC area), a metric based on signal-detection theory, and Rayleigh statistics, respectively.

The present study examines the effect of noise exposure on the vestibular periphery by applying a wavelet transform to single-trial VsEP waveforms in animals exposed to a 110 dB SPL 1.5-kilohertz-centered 3-octave band noise. Our previous study reported a temporary noise-induced VsEP threshold shift by analysis of the amplitudes and latencies of trial-averaged VsEP waveforms (Stewart et al., 2021). Our present study reveals that exposure to 110 dB noise for 6 hours significantly reduces the power/strength of the VsEP waves as well as their temporal coherence across trials 1 day after exposure, which then recovers to near pre-exposure levels within 3-7 days and completely by 28 days on a population level. The noise-induced deficit was largest in response to the smallest jerk stimulus, indicating that an animal’s sensitivity for detection of small jerk head motions was affected by noise exposure. We also found that the number of calretinin-positive (CR+) calyces in the sacculi harvested 28 days after noise exposure was significantly reduced. Furthermore, there was a significant correlation of CR+ calyces with a reduction in temporal coherence across trials and/or signal detectability (ROC area) of the smallest-amplitude jerk. The wavelet transform analysis of the VsEP on a single-trial basis provided more detailed descriptions of noise-induced changes in VsEP waveforms, was a more sensitive tool for assessment of VsEP thresholds, and provided new insights into the neural activity that underlies the VsEP.

## Materials and Methods

### Animals

The VsEP data used in this present study were from 18 out of 21 Long-Evans rats used in Stewart et al. (2021). Remaining three animal’s data were not analyzed due to difference in the file format and not included in the present study. The rats were male and their weight ranged between 350 ∼ 400 g (Charles River Laboratories, MA). They were caged in pairs with free access to food and water. The housing facility had a 12/12-hour light/dark cycle. The rats were subjected to ABR and VsEP measurements before and after a 6-hour long exposure to 110 dB noise. All animals were euthanized on 28 days after exposure, and their saccules were collected for an immunohistochemistry. All procedures were carried out in accordance with National Institutes of Health guidelines and were approved by the Institutional Animal Care and Use Committee at the University of Michigan.

### Experimental design

Each rat was surgically implanted with a small head bolt that was designed to be fixed onto an electromagnetic shaker during VsEP recordings (see Surgical implantation). After post-operative recovery, baseline VsEP was measured (see VsEP recording). Then, the animal was exposed to 110 dB SPL noise for 6 hours (See Noise exposure). The VsEP was measured 1, 3, 7, 14, 21, and 28 days after exposure. Additionally, the ABR was measured before and 1, 3, 7, and 28 days after noise exposure, but those results are not part of the present study.

After the completion of the physiological recordings, animals were euthanized, their saccules were processed for calretinin immuno-staining, and calretinin-immunoreactive calyces were counted (see Immunohistochemistry). Due to technical difficulties during dissection and immuno-staining, images of saccules from 6 animals were not available for further analyses. Thus, the present study reports on the saccular data from a partial set of animals (12 out of 18). We used a normative set of 13 animals without noise-exposure, who were in a similar age range at the time of tissue collection, as controls to examine the change in calretinin-positive calyces following noise exposure.

#### Surgical implantation

Details were described in Stewart et al. (2020, 2021). Briefly, rats were anesthetized with isoflurane and placed on a stereotaxic frame, where a midline incision exposed the bregma and lambda on the dorsal surface of the skull. The stereotaxic positioning was refined so that the bregma and lambda were at equal depths in the z-plane. Two anchor screws were placed into the skull. A custom, small bolt was placed vertically against the bregma and cemented with C&B Metabond (Parkell, NY). Dental acrylic was fused around the bolt and the anchor screws to strengthen the attachment of the head bolt to the skull. The animal was placed on post-operative recovery for 10 days.

#### Noise exposure

Details were described in Stewart et al. (2020, 2021). Briefly, unanesthetized rats in individual wire mesh cages were placed into a ventilated noise exposure booth. They received a free-field, 3-octave noise with a center frequency of 1.5 kHz for 6 hours. The sound level was calibrated with a fast Fourier transform spectrum analyzer (SR760; Stanford Research Systems, CA) by setting the peak of the frequency spectrum to 110 dB SPL.

#### VsEP recording

VsEP was recorded as described previously (Stewart et al., 2020, 2021). Briefly, a rat was anesthetized with a mixture of ketamine and xylazine and its head was affixed to a shaker (ET-132, Labworks Inc., CA) via the surgically implanted bolt. Electrodes (stainless needles) were placed subcutaneously at the vertex (noninverting), mastoid (reference), and hip (ground). Jerk stimuli with 5 different amplitudes (0.32, 1.1, 2.2, 3.2, and 5.5 g/ms) were produced by applying linear voltage ramps of different slopes and durations to the shaker. Jerk is the rate of change in linear acceleration. The direction of the stimulus aligned with the animal’s naso-occipital axis. Jerk stimuli of a single amplitude were delivered as successive positive- and negative-polarities and repeated for 200 repetitions. In each block of recording, 5 jerk levels were tested in an increasing order. Typically, 3 blocks of recordings were performed on each animal on a given day of recording.

#### Immunohistochemistry

The number of calretinin-positive calyces was determined as described previously (Stewart et al., 2020). Briefly, after animals were given a lethal dose of sodium pentobarbital, otic capsules were removed, immersed in the fixative for 4 hours or longer up to overnight, and decalcified in 5% EDTA in PBS for 3∼4 days at room temperature. The utricle and saccule were dissected, then blocked and permeabilized in 0.3% Triton X-100, 5% normal donkey serum in PBS with Ca^2+^ and Mg^2+^ for 2 hours at room temperature. Sacculi were immersed in a combination of primary antibodies against mouse anti-calretinin (CR; 1:1000, Millipore MAB1568), chicken anti-neurofilament H (NF; 1:1,000, Millipore AB5539), and rabbit anti-myosin 7a (Myo7a; 1:100, Invitrogen, PA1-936), followed by respective secondary antibodies. Sacculi were then whole-mounted on slides in ProLong Diamond Antifade Mounting Medium (Invitrogen) and coverslipped. A 40X z-series with 0.3-μm steps were collected with a confocal microscope (Stellaris, Leica, Germany) at a region of interest (Stewart et al., 2018, 2020). Images were analyzed for the presence of CR+ calyces with FIJI software (ImageJ). CR+ calyces are defined as CR-positive, Myo7a-negative, and NF-positive circles of diameter greater than 5 μm that appeared below the cuticular plate of the sensory epithelium.

#### Wavelet transform

The Wavelet Toolbox from MATLAB R2022b (Mathworks, MA) was used in our time-frequency analysis of the VsEP with wavelet transform. We used a continuous Morse wavelet transform (MATLAB function ‘cwt’) with a symmetry parameter, gamma (*γ*), at 3 (symmetrical in time domain), a time-bandwidth product, P, at 10 (the square root of *P* is proportional to the wavelet duration in time), and a redundancy of 10 voices per octave (frequency points per octave in wavelet transform), applied on a 12-ms trace of the single-trial VsEP, encompassing 6 ms before and 6 ms after the onset of a jerk. We did not post-process the voltage recording (e.g., filtering) before applying the wavelet transform. Example wavelets at 3 different center frequencies are shown in Fig. 2D. Because we were interested in discriminating different components of the VsEP (e.g., P1/N1 vs P2/N2), the time-bandwidth product was selected at 10 so as to shorten the temporal duration of the wavelets as much as possible without compromising their frequency selectivity.

**Figure 1.**
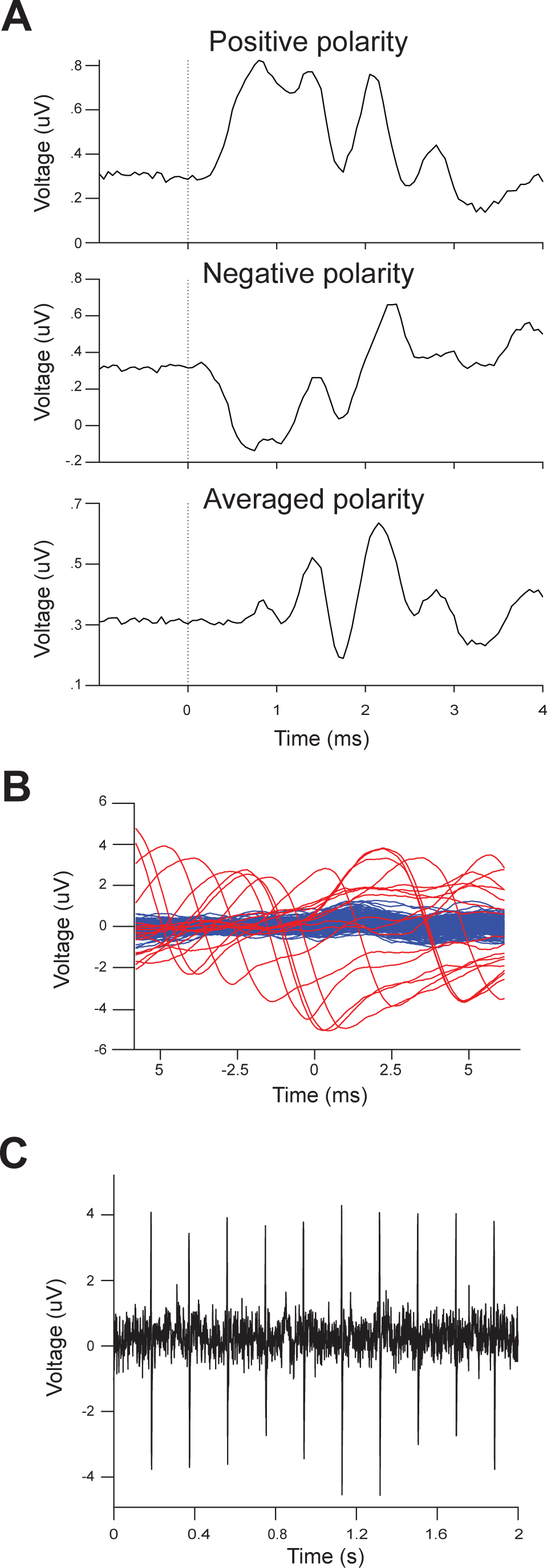
**A**: VsEP traces were generated by presenting 200 paired trials of positive-(top panel) and negative-(middle panel) polarity jerks at 5.5 g/ms and averaging the response of trials that did not show cardio-vascular related noise (red traces in **B**). The response to positive- and negative-jerks were further averaged to produce the polarity-averaged response (bottom panel). **B**: Low-pass filtered voltage recording of the VsEP aligned at the onset of jerks (t = 0) shows slow fluctuations with a stereotypical shape occurring at random timing relative to the onset of jerks, likely cardio-vascular related noise. Trials that include this type of noise (red traces) were removed from our analysis by setting an empirical threshold based on voltage and/or derivative of voltage trace. **C**: A longer segment of the VsEP recording (2 s) shows the large fluctuations in **B** occurring periodically at an interval of ∼0.188 sec, which equates to ∼320 per minutes.

**Figure 2.**
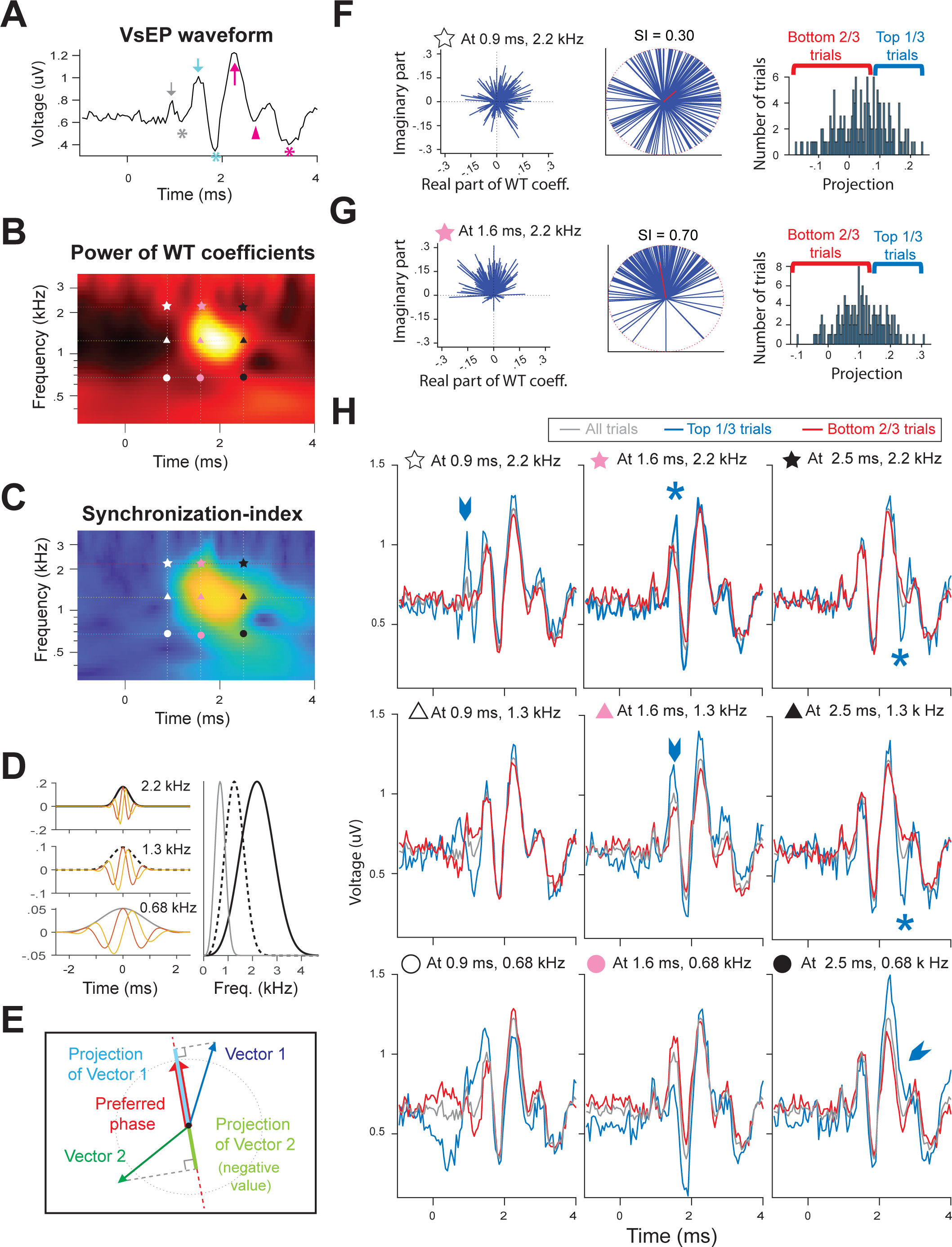
**A**: The VsEP trace generated by averaging 161 paired trials of positive- and negative-polarity jerks at 5.5 g/ms. VsEP peaks P1, N1, P2, N2, P3, and N3 peaks are denoted with a gray arrow, gray asterisk, blue arrow, blue asterisk, red arrow, and red asterisk, respectively. A red arrow-head denotes a N3’ dip. **B**: Wavelet transform was performed on single-trial VsEP waveforms in **A**. The power (absolute value) of the wavelet transform coefficients were averaged across 161 trials and shown as a two-dimensional (Time x Frequency) heat map. **C**: Synchronization-index of phases of the wavelet transform coefficients for the waveform in **A,** showing how well the phase of the wavelet transform coefficients are aligned across trials. In **B** and **C**, star, triangle, and circle symbols in white, pink, and black colors indicate a point where wavelet transform coefficients are taken and examined in **F**∼**H**. **D**: On the left 3 panels, the wavelet kernels are shown for the center frequencies of 2.2 (top), 1.3 (middle), and 0.68 kHz (bottom). The right panel shows frequency response profiles for 2.2 (black solid), 1.3 (black dotted), and 0.68 kHz (gray solid) wavelet kernels. **E**: Schematics of the projection of a given vector onto the preferred-phase vector (red). **F** and **G**: Left panels: Distribution of wavelet transform coefficients (complex numbers) is shown on a complex plane for those taken at 0.9 ms and 2.2 kHz (**F**) and at 1.6 ms and 2.2 kHz (**G**). Middle panels: Distribution of the phase of the wavelet transform coefficients in left panels is shown on a unit circle (red, dotted circle). Each blue, solid line indicates a unit vector representing the phase of a wavelet transform coefficient from a single trial. The vector average of all blue vectors is depicted with a red, solid vector, representing the preferred phase at the given frequency and time point. Synchronization-index (SI) is the length of the preferred phase vector, representing how well the phase are clustered across trials. Right panels: Distribution of the projection of wavelet-transform-coefficient vectors in left panels onto the preferred phase shown in middle panels. **H**: Trials were sorted according to the projection at a given frequency and time point (indicated at the top of each plot) and grouped into the top one-third and the bottom two-third trials. Average waveforms for the top one-third trials (blue), the bottom two-third trials (red), and all trials (grey) are compared.

#### Synchronization-index and Rayleigh statistics

The Synchronization-index, also known as Vector Strength, was determined as:

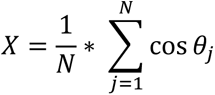

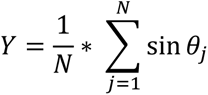

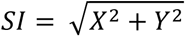

where θ_j_ is the phase of the WT coefficient (determined with the MATLAB function ‘angle’) for the j-th single-trial trace of the VsEP and N is the number of trials. Rayleigh statistics were then determined as:

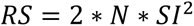

where N is the number of trials. An SI giving rise to RS above 13.816 was considered to indicate a significantly non-uniform distribution of the phases of WT coefficients at p < 0.001 (Mardia and Jupp, 2000).

In order to determine the minimum stimulus level with a significant synchronization in the phase of WT coefficients, the SI was first averaged across three frequency regions (0.54∼0.92, 0.92∼1.58, and 1.58∼2.7 kHz), and then averaged across recording blocks. Then RS was calculated with the average number of trials across blocks as N. The phases of WT coefficients were judged significantly synchronized if RS exceeded 13.816 within a time window of 0 ∼ 4 ms.

#### ROC area calculation

The power of the wavelet transform (WT) coefficients after the onset of jerk stimulation, which represents the ‘signal’ response, was averaged in a time-window (e.g., 0.6 ∼ 3.8 ms) over a range of frequencies (e.g., 1.58∼2.7 kHz) for each trial. Similarly, the power of WT coefficients prior to the onset of jerk (−0.3 ∼ −0.1 ms), which represents the ‘noise’ or baseline response, was averaged over the same frequency range for each trial. From these 2 distributions, we calculated (1) the true positive rate, the probability of correct detection of a jerk, and (2) the false positive rate, the probability of incorrectly reporting the presence of a jerk (false alarm) by use of a decision criterion selected from all experimentally obtained values of the power (including both the signal and noise response). The receiver operating characteristics (ROC) curve is a plot of the true positive rates against the false positive rates (Green and Swets, 1988). The area under the ROC curve, ROC area, ranges from 0 to 1. An ROC area of 1 or 0 means that an ideal observer can perfectly detect the presence of a signal (the jerk) based on an increase or decrease in the power of the WT coefficients, respectively. An ROC area of 0.5 means chance detectability.

## Statistical Analysis

We used built-in functions of MATLAB R2022b (Mathworks, MA) to perform paired t-tests (‘ttest’), two-sample t-tests (‘ttest2’), signed-rank tests (‘signrank’), and Pearson’s correlation tests (‘corr’ with type = ‘Pearson’). Additionally, a custom program was developed in MATLAB to test for the significance of ROC area using a permutation test (Britten et al., 1996). First, the power of WT coefficients from ‘signal’ and ‘noise’ responses were pooled together. Random samples were taken from the pool, without replacement, and assigned to ‘signal’ and ‘noise’ groups in an equal proportion. The ROC area was then calculated from these 2 distributions of random samples. This procedure was repeated 1000 times. The p-value < 0.01 means that less than 1 % (10 out of 1000) of the randomization process produced an ROC area greater than the experimentally obtained ROC area.

## Results

The vestibular short-latency evoked potential (VsEP) is a far-field compound action potential evoked by a ramp of linear acceleration applied to the animal’s head. The rate of change in linear acceleration is termed “jerk”, and the amplitude of the VsEP has been shown to primarily depend on this parameter (Jones et al., 2011). Experimentally, a VsEP waveform is obtained by presenting a pair of positive and negative jerks to an animal for multiple repetitions and averaging the responses to both polarities. Positive and negative jerks produce stimulus-locked waves in opposite directions, and averaging them cancels out the “artifact” thought to be generated by the electro-magnetic shaker that produces the jerk stimulus (Fig. 1A). In addition, a large fluctuation in voltage, likely of cardiovascular origin, was present in our recording (Figure 1B and 1C). The noise was highly periodic (but unsynchronized with the jerk stimuli), and its rate (∼320 per minute in the example of Fig. 1C) is at the lower end of the range of the heart rate reported for awake, undisturbed rats (between 300 to 425 beats per minutes; Azar et al., 2011). We removed trials that contained this type of noise from our analysis (red traces in Fig.1B).

The typical VsEP trace shown in Fig.2A was obtained by presenting 200 trials of positive and negative jerks at 5.5 g/ms and averaging the response from 161 paired trials after removing 39 trials containing cardiovascular noise. The VsEP typically starts with a positive peak (P1, denoted with a gray arrow), then a negative peak (N1, a gray asterisk), both of which are thought to represent the compound action potentials of irregular afferents in the otolith organs (Nazareth and Jones, 1998). The subsequent P2, N2, P3, and N3 peaks (cyan arrow, cyan asterisk, red arrow, and red asterisk, respectively) are thought to represent the activity of vestibular neurons (either vestibular afferents or their targets in the brain stem; Jones & Jones, 1999; Nazareth & Jones, 1998). Between P3 and N3 peaks, there is a dip, N3’, (denoted with a red arrow head) that is more pronounced in some animals and less so in others.

A wavelet transform was applied to the single-trial VsEP traces after pairs of positive- and negative-polarity jerk responses were averaged. The wavelet transform produces a two-dimensional array of complex numbers, representing the power and the phase of waves of various frequencies over time. The distribution of wavelet transform coefficients across trials is plotted on a complex plane for the coefficients at 0.9 ms and 2.2 kHz (left panel on Fig 2F) and for those at 1.6 ms and 2.2 kHz (left panel on Fig. 2G). Each blue line represents a coefficient from a single trial and its length equals its power. The power of wavelet transform coefficients (WT coefficients) averaged across trials is shown as a heat map in Fig.2B. A clear peak in the power is seen around 1.5∼2.2 ms and 1∼1.4 kHz, corresponding to the large down-stroke from P2 to N2 and then the upstroke to the P3 peak in the waveform (Fig.2A).

In addition to the information regarding the power of a wave at a given frequency and time, the wavelet transform provides information about the timing of these waves across trials. On the left panel of Fig 2F, the angle that each blue line makes with the positive x-axis represents the phase of the WT coefficient. The distribution of the coefficient phases is replotted on a unit circle (red, dotted circle) on the middle panel of Fig.2F by disregarding the power of the coefficients. The vector average of all these unit vectors is depicted with a red line, representing the preferred phase of the coefficients at a given frequency-time point. A Synchronization-index, also known as Vector Strength or Phase Clustering, is defined as the length of the resulting preferred phase vector (Cohen, 2014). The synchronization-index (SI) represents the consistency of coefficient phases across trials, ranging from 0 (uniform distribution) to 1 (perfect synchronization). Rayleigh statistics were used to test for statistically non-uniform distributions of phase and indicated that the phase distribution in Fig.2F (and also 2G) is significantly clustered (p<0.001; see Synchronization-Index and Rayleigh statistics in Materials and Methods). The SI of WT coefficients is shown as a heat map in Fig.2C. A peak in the SI map roughly co-localizes with the peak of the power map (Fig.2B), indicating that the wave with 1∼1.4 kHz frequency around 1.5∼2 ms, is temporally best aligned across trials in addition to having the largest power.

One drawback of the time-frequency analysis using a wavelet transform is that we lose temporal precision in exchange for information about frequency. The three left panels on Fig.2D show 3 example wavelets in the time domain, demonstrating, on the one hand, that the temporal duration of these wavelets is greater for the wavelets with lower center frequencies. On the other hand, the frequency response profiles of these 3 wavelets (right panel in Fig.2D) are wider for higher center frequencies, indicating that the frequency selectivity is better for wavelets with lower center frequencies. Thus, there is always an uncertainty in temporally (and spectrally) localizing WT coefficients, and the loss in temporal precision is greater for lower frequencies.

To help understand the relationship between the VsEP waveform and WT coefficients, we sorted trials based on the power and timing information provided by the wavelet transform and then examined the waveform produced by the sorted partial sets of trials. First, a WT coefficient from a single trial at a given frequency-time point was projected onto the preferred phase, treating both as vectors (Fig.2E). The projection of this coefficient relative to the preferred phase vector takes into account both the power and timing information of the WT coefficients; the greater the power and the better the timing with respect to the preferred phase vector, the larger the projection onto the preferred phase vector. Trials were sorted into two groups by ranking them by their projections (right panels in Fig.2F and 2G): (1) top one-third group and (2) bottom two-thirds group.

The average VsEP trace obtained by averaging the top 1/3 of trials selected from their projections at 0.9 ms and 2.2 kHz enhanced the P1-N1 component in the VsEP waveform (blue trace in top, left panel in Fig.2H) compared to a VsEP waveform averaged across all trials (grey trace). In contrast, the VsEP waveform obtained by averaging the bottom 2/3 of trials produced a much-reduced P1-N1 component (red trace). This indicates that the WT coefficients around 0.9 ms and 2.2 kHz are related to the P1-N1 component of the VsEP waveform.

For the P2-N2 component of the VsEP, sorting and averaging trials based on the projections at 1.6 ms and 1.3 kHz generated an enhanced P2-N2 peak compared to the waveform averaged across all trials (blue arrow head on a center panel in Fig.2H). Trials sorted on projections at the same time point (1.6 ms), but at the higher center frequency of 2.2 kHz (top middle panel) also enhanced a P2-N2 peak, but with a sharper down-stroke and a more pointed P2 peak. The subtle difference in the shape of the P2-N2 wave may be due to the broad frequency selectivity of the high-frequency wavelets.

For the P3-N3 component, sorted trials based on the projections at 2.5 ms and 0.68 kHz produced an enhanced P3-N3 component without a prominent N3’ dip on the down-stroke when the top 1/3 of trials were selected (blue arrowhead in the bottom, right panel in Fig.2H). Trials sorted on projections at the same time point (2.5 ms), but at the higher center frequency of 1.3 kHz or 2.2 kHz (right panels on the top and middle rows in Fig.2H) produced a sharp downstroke from P3, creating an obvious N3’ dip.

This analysis demonstrates that WT coefficients at different time points and different frequency bands can be related to different components of the VsEP waveform, although an uncertainty in both time and frequency domains exists.

### Effect of a single exposure to 110 dB noise on the timing of VsEP waves

A representative animal’s VsEP waveforms are shown in Fig. 3A for 0.32, 1.1, 2.2, 3.2, and 5.5 g/ms jerk levels (stimulus level 1, 2, 3, 4, and 5, respectively, from left to right panels). The VsEP was measured prior to noise exposure (black solid traces on top row), 1 day (red), 3 days (cyan), 7 days (magenta), and 28 days (green) after noise exposure. The Synchronization-index (SI) of WT coefficients on the single-trial VsEP waveforms in Fig.3A is shown as two-dimensional (Time x Frequency) heat maps in Fig.3B. Clearly, SI was reduced 1 day after noise exposure for all stimulus levels, indicating that trial-by-trial jitter (i.e., temporal coherence) of VsEP responses was worsened immediately after noise exposure. It then substantially recovered within 3∼7 days.

**Figure 3.**
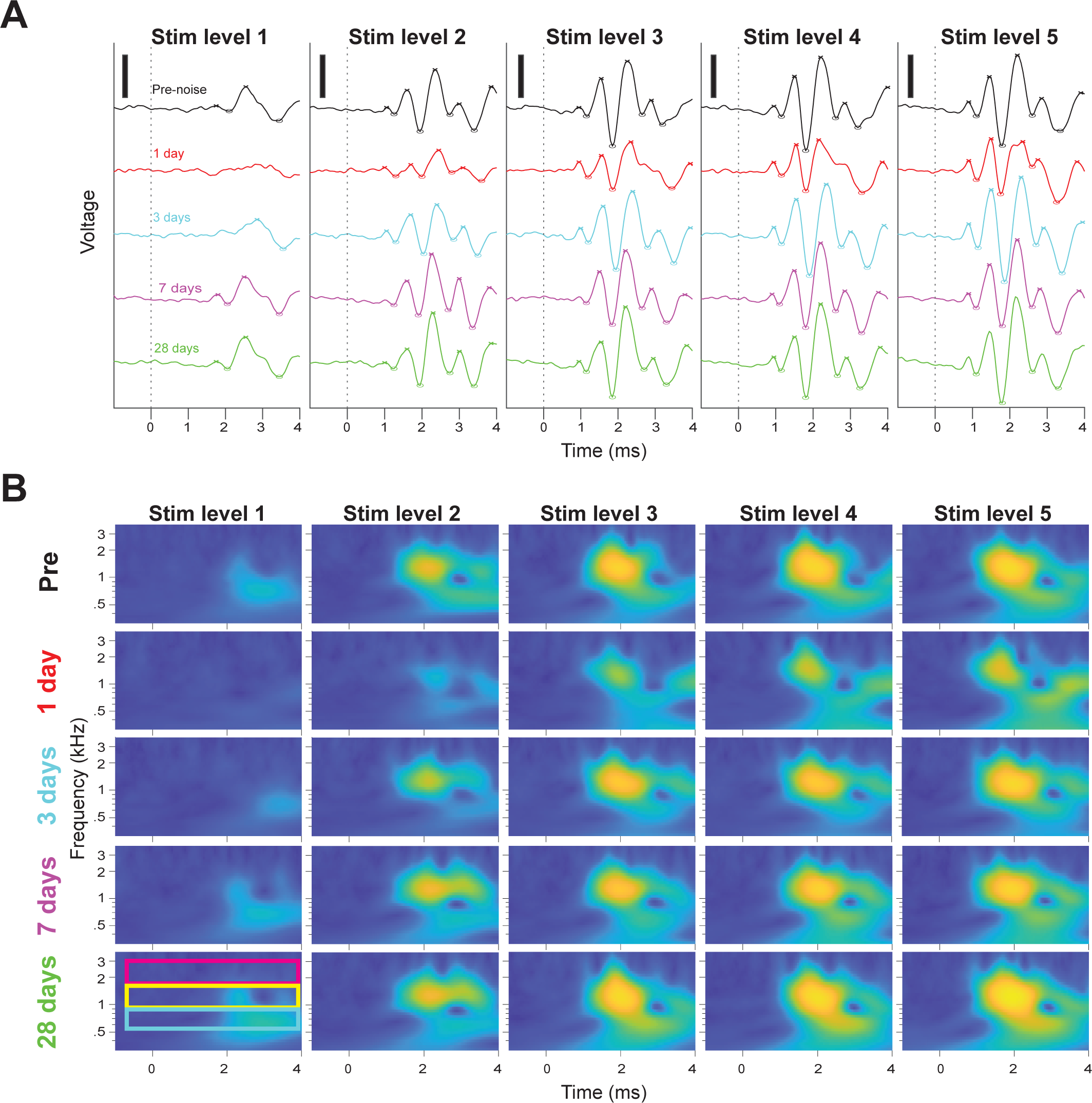
**A**: VsEP waveforms in response to stimulus levels 1∼5 (0.32, 1.1, 2.2, 3.2, and 5.5 g/ms, respectively, from left to right panels) measured before a single exposure 110 dB noise (black solid traces on top row), 1 day (red), 3 days (cyan), 7 days (magenta), and 28 days (green) after exposure from a representative animal. **B**: Synchronization-index (SI) of WT-coefficient phases obtained from the single-trial VsEP waveforms on **A** is shown as a two-dimensional (Time x Frequency) heat map for stim level 1 through 5 (from left to right columns) at pre-exposure (top row), 1 day (2^nd^ row), 3 days (3^rd^ row), 7 days (4^th^ row), and 28 days (bottom row) after noise exposure.

To be better able to quantify the effect of noise exposure, SI was averaged across 0.54 ∼ 0.92 kHz (range depicted with a blue box in the bottom, left panel of Fig.3**A**), 0.92 ∼1.58 kHz (yellow box), and 1.58 ∼ 2.7kHz (red box). These frequency ranges have an equal spectral width in a log scale (which equalizes differences in the spectral selectivity between wavelets with different center frequencies) and are also set around the frequencies shown in Fig.2H, in order to demonstrate the effect of noise exposure on the P1-N1, P2-N2, and P3-N3 components of VsEP. Average SI in high, middle, and low frequency regions is plotted as a function of time in Fig.4A to compare pre-exposure values (black) to 1 day (red), 3 days (cyan), 7 days (magenta), and 28 days (green) after noise exposure. Horizontal dotted lines represent the threshold level of significant synchronization based on Rayleigh statistics (see Synchronization-index and Rayleigh statistics in Materials and Methods).

**Figure 4.**
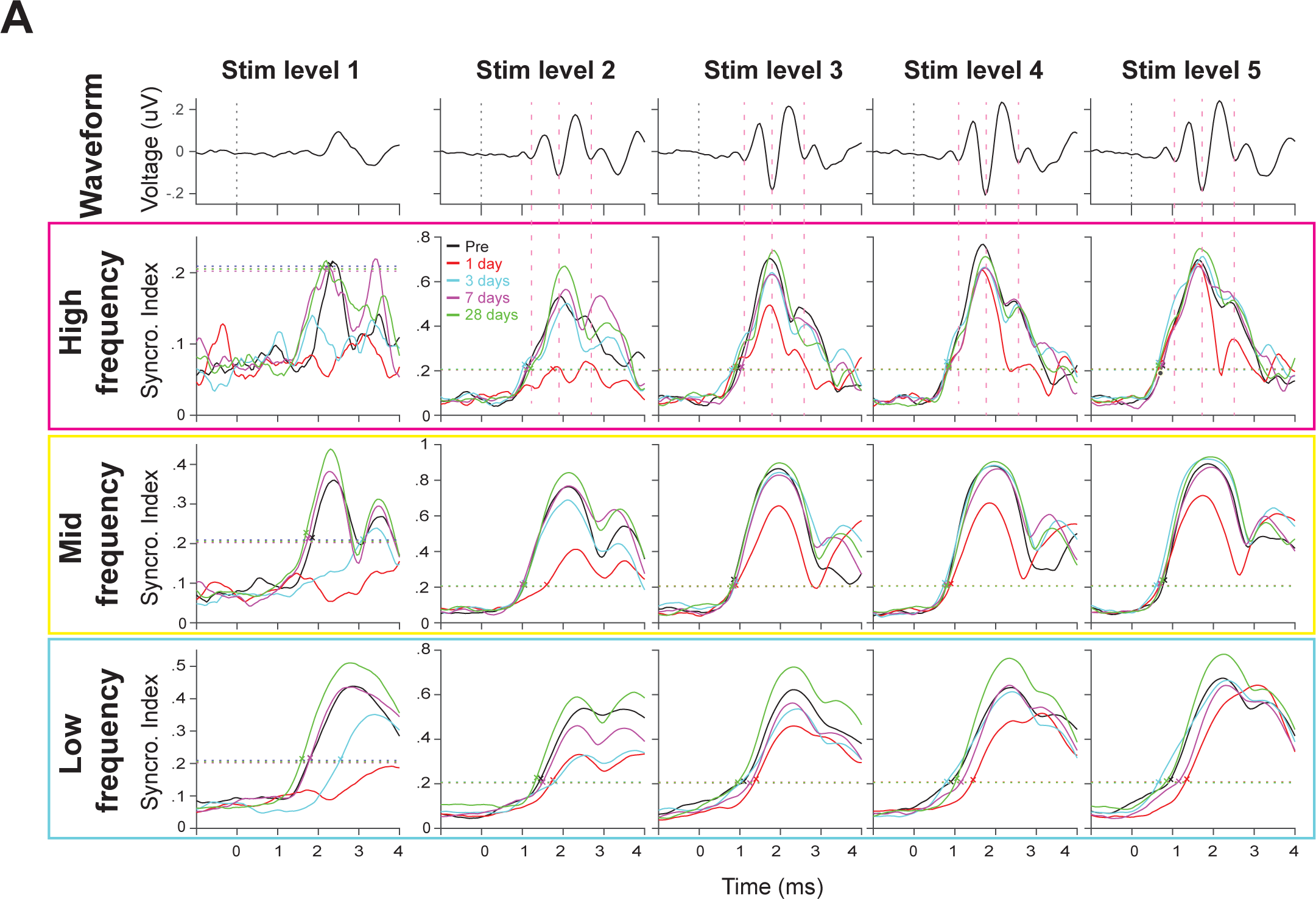
**A**: Synchronization-index is averaged across 0.54∼0.92 kHz (range depicted with a blue box in the bottom-left panel of Fig.3B), 0.92 ∼1.58 kHz (yellow box), and 1.58∼2.7kHz (red box) and plotted as a function of time to compare pre-exposure values (black) to 1 day (red), 3 days (cyan), 7 days (magenta), and 28 days (green) after exposure. Stimulus level 1 through 5 are shown from left to right columns. Horizontal dotted lines indicate the threshold level of significantly non-uniform SI based on Rayleigh statistics at p < 0.001 level. Pre-exposure VsEP waveforms are shown in the top row panels for stim level 1 to 5 (from left to right), and the dotted, magenta lines indicate the times of N1, N2, and N3’-dip.

The SI in the high frequency region, where the greatest temporal precision is expected due to the shortest duration of the wavelets, exhibits a temporal structure with a clear peak that roughly corresponds to the N2 peak in the VsEP waveform (panels in top row of Fig.4A). There is a secondary peak in the high-frequency SI that roughly corresponds to the VsEP N3’ dip. There is no clear peak in the high-frequency SI corresponding to the N1 peak, although an inflection point in the rise of SI from the onset of stimulus to the peak roughly corresponds to the time of N1 peak. Also, SI reaches above threshold level on/before the time of N1 peak for all but the lowest stimulus level (except at 1 day after exposure), indicating that there is significant phase-coherence across trials during the time of P1-N1 component.

At stimulus levels 4 and 5, the high-frequency SI 1 day after noise exposure shows the greatest reduction around the time point when the P3 peak and N3’ dip occur (two right panels on the 2^nd^ row of Fig.4A). This is consistent with less prominent N3’ dips and blunted P3 peaks in the corresponding VsEP waveforms (2^nd^ row in Fig.3A). At lower stimulus levels 1∼2, the SI on day 1 is greatly affected for the earlier time points (two left panels on the 2^nd^ row of Fig.4A). High-frequency SI reaches a level of significance (horizontal, black dotted line) at stimulus level 1 on pre-exposure, 7 days, and 28 days after noise exposure (top left most panel in Fig.4A); on 1 and 3 days after noise exposure, it reaches significance only at stimulus level 2 and above (top, 2^nd^ from the left most panel).

For the middle and low frequency regions, SI is greatly reduced 1 and 3 days after noise exposure for the lowest stimulus level. For stimulus levels 3∼5, the degree of reduction in SI is smaller and quickly recovers by day 3. The minimum stimulus level with significant SI exhibits a temporary threshold shift from stimulus level 1 before exposure to stimulus level 2 at day 1 of exposure for both mid and low frequency regions, then returns back to stimulus level 1 at day 3. This pattern matches well with the thresholds determined by examination of trial-averaged VsEP waveforms.

As a population, SI was significantly reduced 1 day after noise exposure for high, mid, and low frequency ranges, but largely recovered to pre-noise level within 3∼7 days of noise exposure (Fig.5A). In Figure 5A, time windows that have significant difference in SI between pre- vs post-exposure (by paired t-test at p < 0.05) are indicated by horizontal bars in the same color as corresponding population-mean SI traces. At the smallest jerk level, the decline in SI in high and mid frequency regions gradually recovered from day 1 (red) to day 3 (cyan), day 7 (magenta), then day 28 (green). Note that there was individual variability such that some animals did not recover from damage, while others completely recovered or even showed an increase in SI, resulting in a population average 28 days after noise exposure that was indistinguishable from the pre-exposure level.

**Figure 5.**
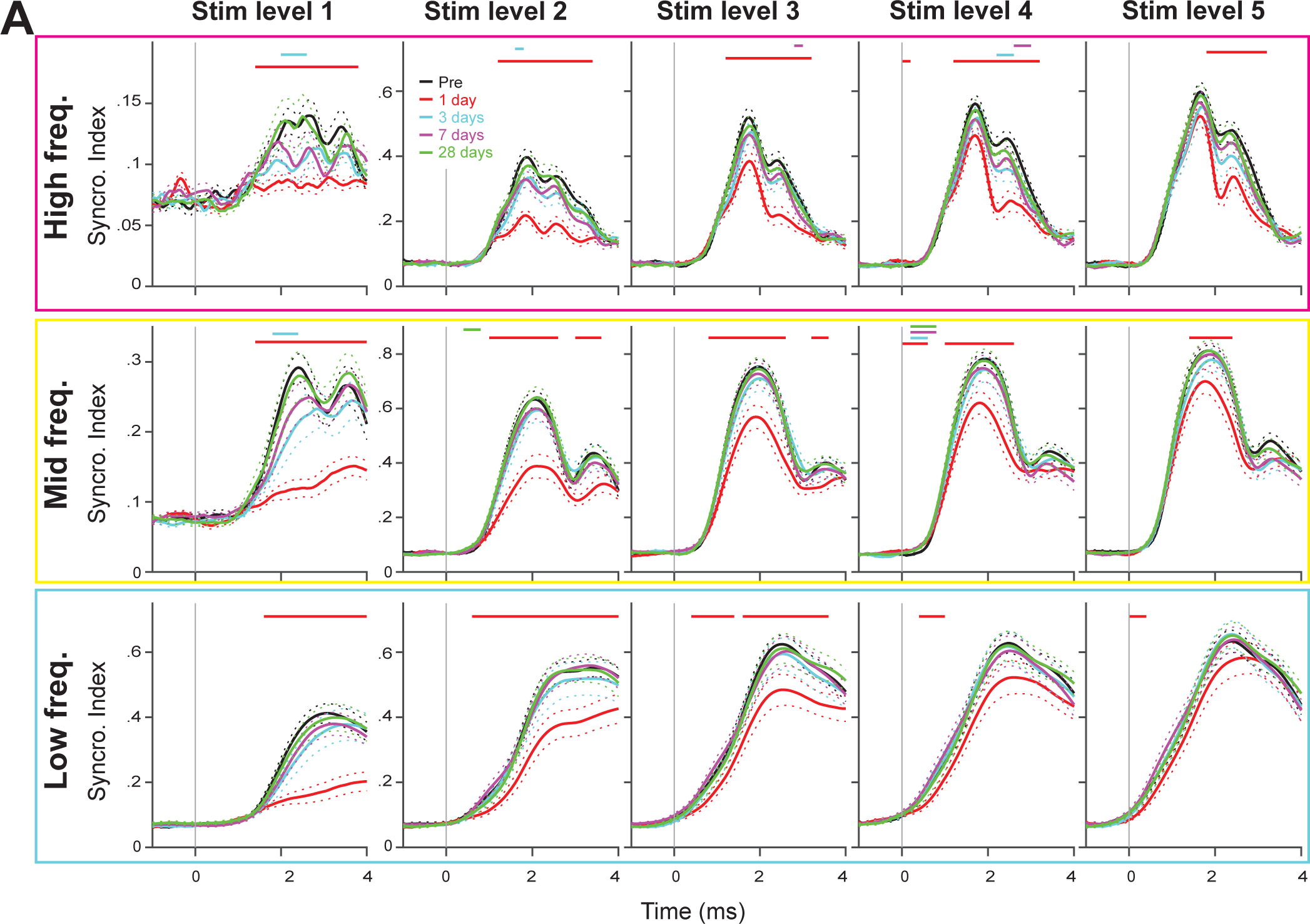
**A**: Population-average Synchronization-index (SI) in high (1.58∼2.7 kHz, top row), middle (0.92 ∼1.58 kHz, middle row), and low frequency regions (0.54∼0.92 kHz, bottom row) for stimulus level 1 through 5 (from left to right columns). Pre-exposure (black), 1 day (red), 3 days (cyan), 7 days (magenta), and 28 days (green) after noise exposure. Dashed traces indicate standard errors in mean. Horizontal, solid bars at the top of each panel represent the time range, where the post-exposure mean SI is significantly different from pre-exposure with paired t-test at p = 0.05 level. The color of the bar denotes the exposure days that had significant difference (red: 1 day, cyan: 3 days, magenta: 7 days, green: 28 days). The test was performed by comparing SI averaged within a 0.2 ms time bin between two conditions.

At stimulus level 5, high-frequency SI 1 day after exposure showed the greatest reduction around a later time period corresponding to P3-N3’ component, while the earliest part did not show significant decline compared to pre-exposure. However, at lower stimulus levels, the SI reduction at 1 day was large across the entire analyzed time frame. For SI in mid and low frequency ranges, a significant reduction was seen 1 day after noise exposure, but the effect was again greater for lower stimulus levels.

Reflecting the immediate effect on SI, which was greatest at the lowest jerk level, the majority of animals showed an increase in the minimum stimulus level where SI reached a significant level (termed threshold stimulus level), one day after exposure. For high-, mid-, and low-frequency regions, threshold stimulus level increased in 44.4%, 72.2%, and 61.1% of animals, respectively, one day after exposure (Fig. 6A). In addition, we saw a significant increase in the time point when SI crossed a significant level 1 day after exposure for all frequency regions (Fig.6B), indicating SI was affected even in those animals that did not show changes in threshold stimulus level.

**Figure 6.**
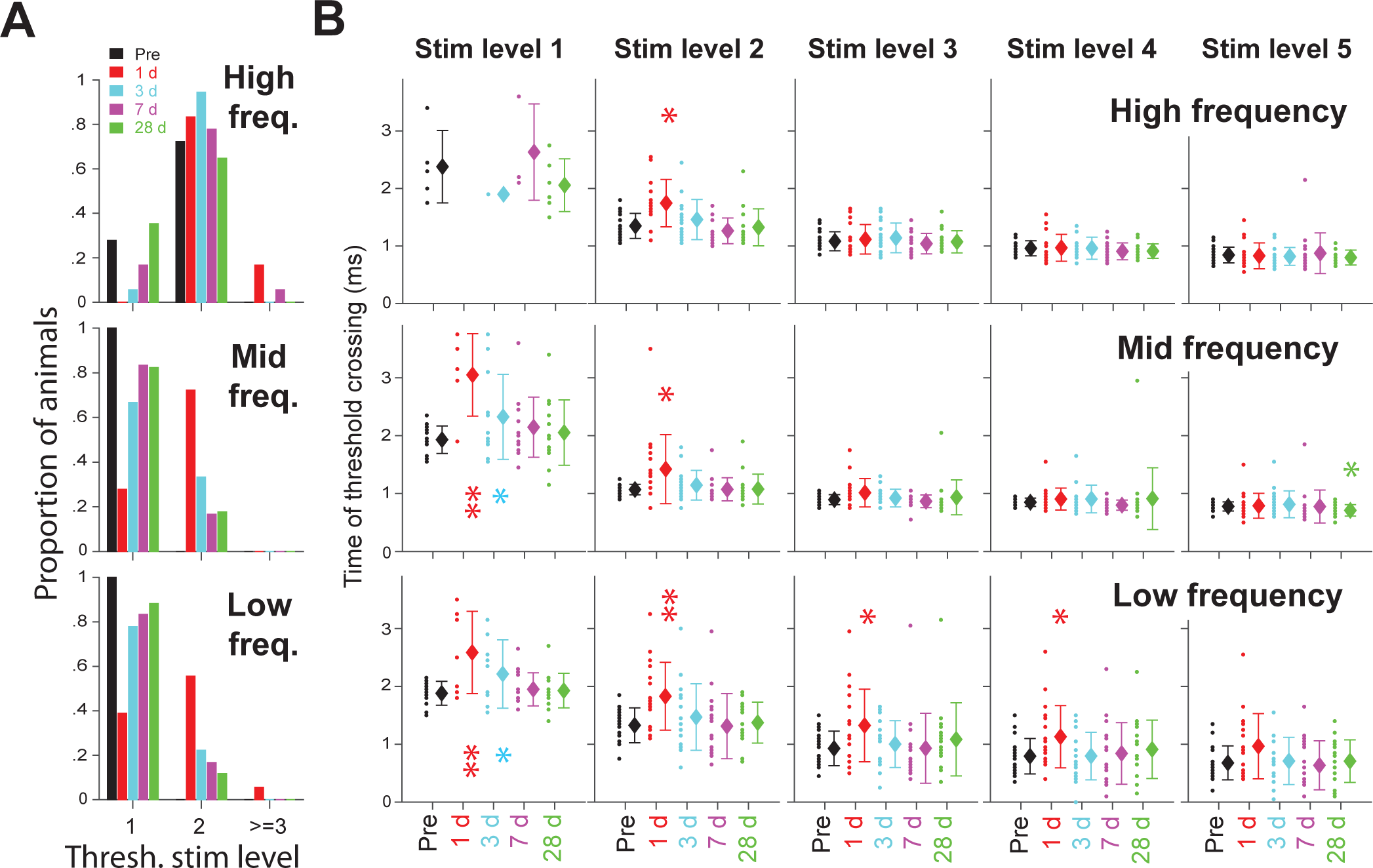
**A**: Distribution of minimum stimulus level where Synchrorization-index (SI) reached a threshold (∼ 0.21 depending on the number of trials, N) before 4 ms of the stimulus onset. **B**: Distribution of time when SI reached a threshold is compared between pre-exposure and 1, 3, 7, and 28 days after noise for stimulus levels 1 through 5 (from left to right columns). Single- and double-asterisks indicate significantly different mean values compared to pre-exposure with two-sample t-test at p < 0.05 and <0.01, respectively. For **A** and **B**, threshold based on SI in high, middle, and low frequency ranges are shown on the top, middle, and bottom rows, respectively.

The threshold stimulus level did not recover to pre-noise levels for at least one frequency region in a minority of animals after 28 days. Five out of 17 (29.4%) animals did not recover in one frequency region but recovered in the other two. Two out of 17 animals only recovered in one frequency region. All animals showed recovery in at least one frequency region.

### Effect of a single exposure to 110 dB noise on the power of VsEP waves

The power of WT coefficients obtained from single-trial VsEP waveforms (the same animal as Fig.3) is shown as a heat map for stimulus level 1 through 5 (Fig. 7A from left to right columns) on pre-exposure (top row), 1 day (2^nd^ row), 3 days (3^rd^ row), 7 days (4^th^ row), and 28 days (bottom row) after noise exposure. Compared to SI maps from the same animal (Fig.3B), the power maps are noisier at the highest and lower frequency regions. Nonetheless, we can see a clear reduction in the power of WT coefficients 1 day after exposure, which recovered over time similar to SI.

**Figure 7.**
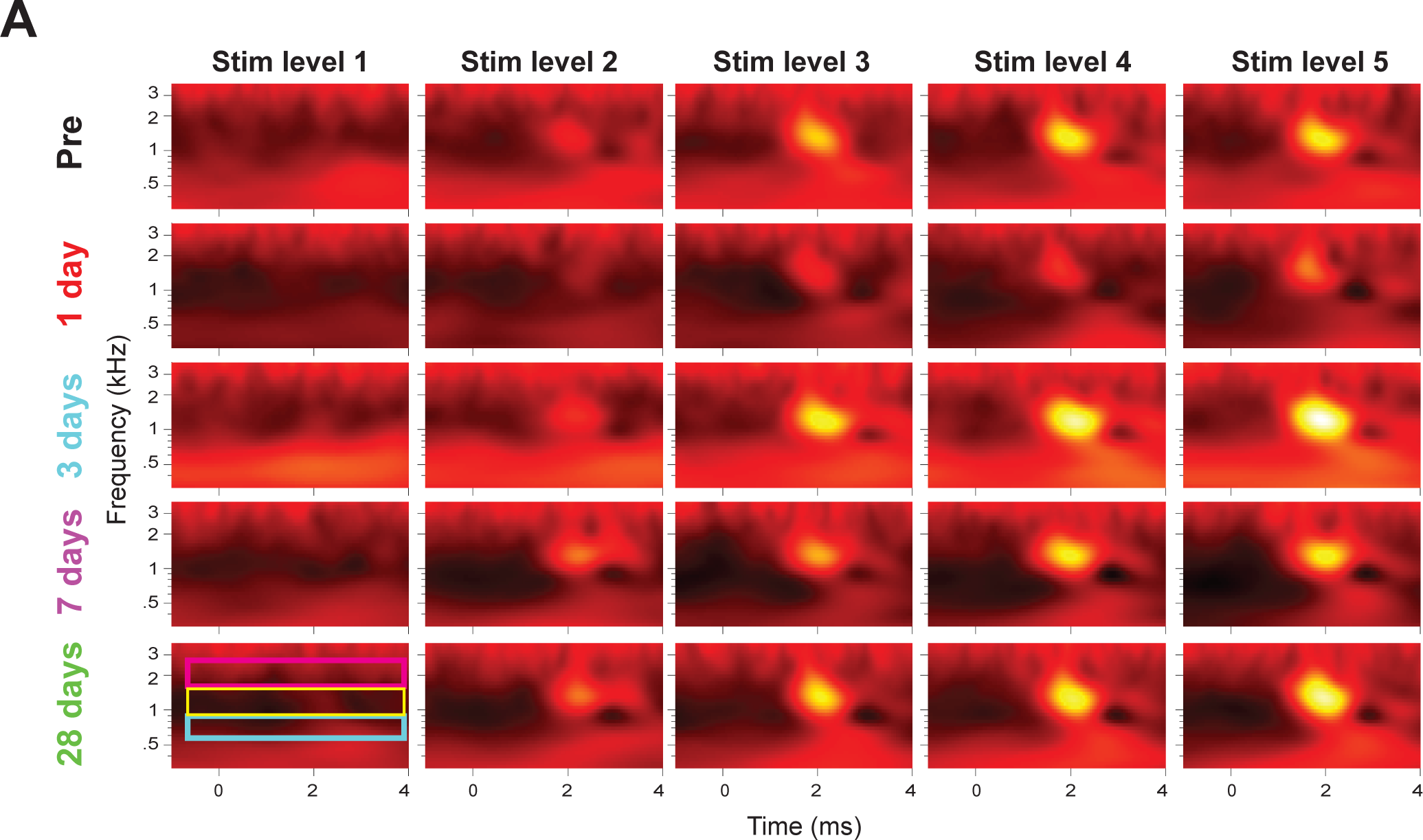
**A**: Power of the wavelet transform coefficients obtained from the single-trial VsEP waveforms on Fig.3A is shown as a two-dimensional (Time x Frequency) heat map for stimulus level 1 through 5 (from left to right columns) at pre-exposure (top row), 1 day (2^nd^ row), 3 days (3^rd^ row), 7 days (4^th^ row), and 28 days (bottom row) after noise exposure.

To quantify the effect of noise exposure, the power of WT coefficients was averaged across the same frequency regions employed for SI analysis (0.54∼0.92 kHz, 0.92 ∼1.58 kHz, and 1.58∼2.7 kHz). In addition, the power averaged across these frequency regions was normalized relative to the ‘background noise’ level calculated in a time window prior to the onset of the stimulus (−0.3 ∼ −0.1 ms) and over the same frequency region. Normalization was necessary because we observed fluctuations in the background noise level between days of exposure (compare the power level prior to time = 0 between days in Fig 7A). The average normalized power in high, middle, and low frequency regions is plotted as a function of time on Fig.8A to compare pre-exposure values (black) to 1 day (red), 3 days (cyan), 7 days (magenta), and 28 days (green) after noise exposure. Black dotted traces show the SI on the pre-exposure day from the same animal in order to show that both SI and the power of WT coefficients peak near a time point that roughly corresponds to the time of the N2 peak in the VsEP waveform. However, SI is more sensitive and rises earlier in time compared to the power of WT coefficients; thus, phase synchronization better represents the P1-N1 component of the VsEP waveform. Overall, the power of the WT coefficients is reduced greatly on day 1 but recovers (or may become larger compared to baseline) 28 days after exposure for all frequency ranges in this representative animal.

**Figure 8:**
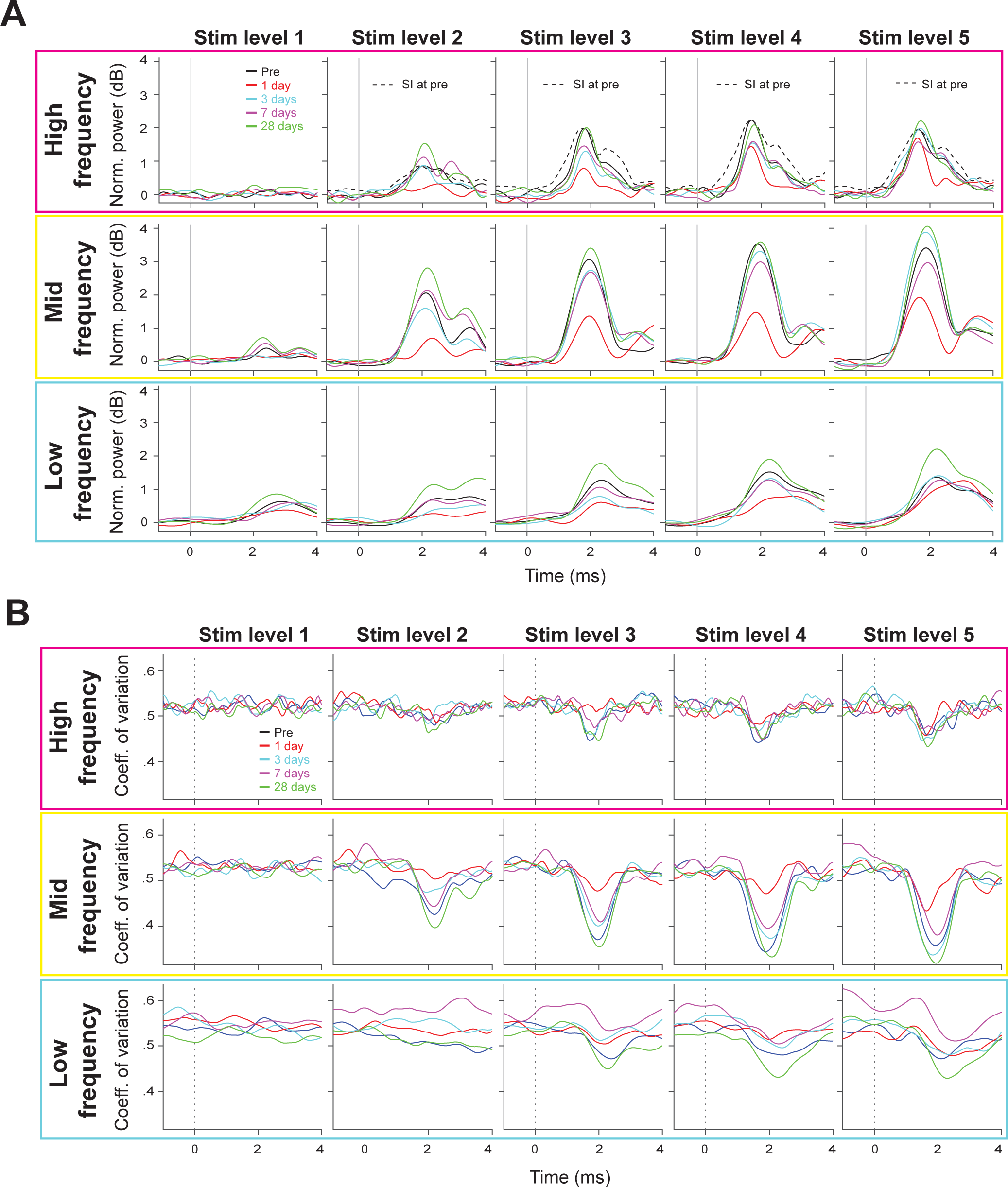
**A**: Normalized power of the wavelet transform coefficients in Fig. 7A is averaged across 0.54∼0.92 kHz (range depicted with a blue box in the bottom, left panel of 7**A**), 0.92 ∼1.58 kHz (yellow box), and 1.58∼2.7kHz (red box) and plotted as a function of time to compare pre-exposure values (black) to 1 day (red), 3 days (cyan), 7 days (magenta), and 28 days (green) after noise exposure. Stimulus level 1 through 5 are shown from left to right columns. Synchronization-index before the noise-exposure from the same animal is shown in black, dotted line in the high-frequency power plots for stimulus level 2∼5 to guide a comparison between the onset of SI and power rising over time. **B**: Coefficient of variation in the single-trial power of the wavelet transform coefficients for the plots shown in **A**.

We next examined trial-by-trial variability in the power. The coefficient of variation (standard deviation divided by mean) in the WT power at high, mid, and low frequency regions is plotted as a function of time on Fig.8B to compare between exposure days. The coefficient of variation decreases after the onset of stimulation, indicating that the jerk-induced response power exhibits less variability compared to the background activity. One day after exposure, the stimulus-induced reduction in variability was smaller, showing that the variability of the response power increased one day after exposure compared to pre-exposure. However, the coefficient of variation quickly recovered by 28 days.

As a population, the mean normalized power of WT coefficients was significantly reduced 1 day after noise exposure for high, mid, and low frequency ranges. However, the population average was indistinguishable from baseline levels 28 days after noise exposure (Fig.9A), although there was individual variability in the degree of recovery. The effect of noise exposure on WT power appeared to be greater for the smallest stimulus levels as was seen with phase-synchronization of the WT coefficients. In the high-frequency region, the normalized power was significantly reduced during the time of the P2-N2 peak to the P3-N3’-N3 peak for all stimulus levels one day after of exposure (see red horizontal bars at the top of Fig. 9A indicating the time period with significant difference at p < 0.05 by paired t-test). This differs with what we observed with SI, where the noise-induced reduction was greatest around ∼2.5 ms (corresponding to P3-N3’ dip components) at stimulus levels 4 and 5.

**Figure 9.**
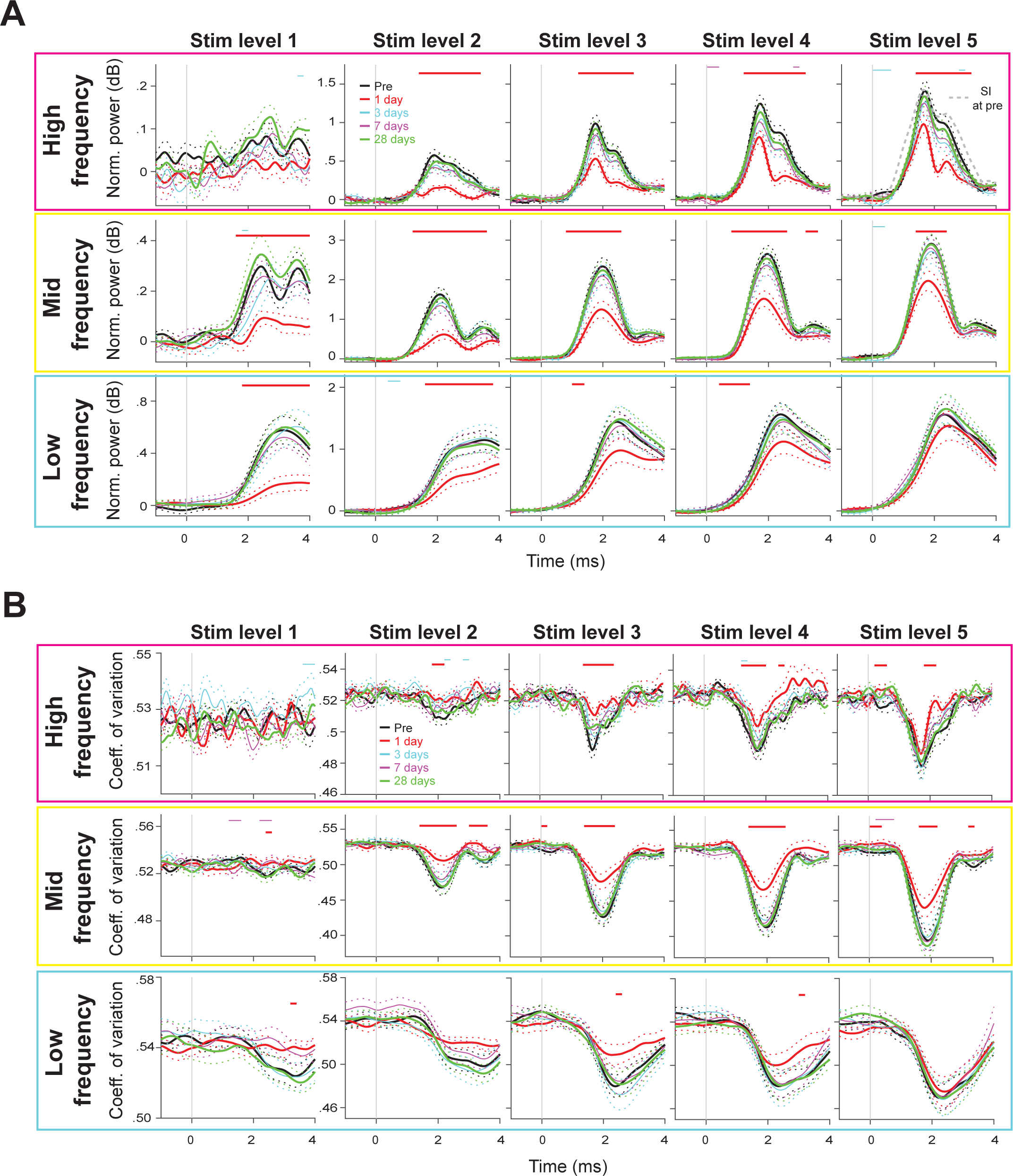
**A**: Population-average normalized power of the wavelet transform coefficients in high (1.58∼2.7kHz, top row), middle (0.92 ∼1.58 kHz, middle row), and low frequency range (0.54∼0.92 kHz, bottom row) for stimulus level 1 through 5 (from left to right columns). Horizontal bars at the top of each panel represent the time range where the post-exposure values are significantly different from the pre-exposure values with paired t-test at p = 0.05 level. The test was performed by comparing the averaged SI within a 0.2 ms time bin between conditions. The population-average SI before the noise-exposure is shown in grey, dotted line in the high-frequency power plots for stimulus level 5. **B**: Coefficient of variation in the single-trial power of the wavelet transform coefficients for the plots shown in **A**.

A comparison of <u>high-frequency</u> SI vs normalized power showed that SI is, on average, more sensitive for detection of the smallest jerks compared to the power of WT coefficients, which appear flat even before noise exposure (top left panel of Fig.9A). In addition, the population-average SI (black dotted trace in top right panel) increases earlier compared to the population-average power of WT coefficients, suggesting that the P1-N1 component of the VsEP is better represented in the timing aspect of the underlying high-frequency waves than in their power.

We also found that the population-averaged coefficient of variation showed a significant increase (manifested as a smaller reduction re background) 1 day after exposure that quickly recovered on subsequent days (Fig.9B). Thus, both the mean and variability in the power of VsEP waves showed temporary impairment that improved substantially within 3 days after noise and recovered to baseline by 28 days after noise on a population level.

Next, we determined the threshold stimulus level of the VsEP based on the power of WT coefficients using a signal detection theory-based metric, the ROC area. ROC area is the area under a receiver operating characteristics curve and represents how well an ideal observer can detect a signal (the occurrence of a jerk) based on the power of WT coefficients. First, the power of WT coefficients averaged over a region of interest on the power map (cyan box in Fig.10A, mid frequency from 0.6 ∼ 3.8 ms) was obtained for each trial (cyan bars in the left panel Fig.10B) and compared to the power averaged over the corresponding frequency region prior to the onset of jerks (−0.3 ∼ −0.1 ms, white box in Fig.10A, its distribution shown in black bars in the left panel of Fig. 10B). From these 2 distributions, an ROC curve was generated by selecting a decision criterion traversing all experimentally obtained values and calculating true and false positive rates (right panel in Fig.10B; Green & Swets, 1988; see ROC area calculation in Materials and Methods). ROC area ranges from 0 to 1; an ROC area of 1 or 0 means a perfect detectability with increased or decreased power, while an ROC area of 0.5 means a chance detectability. A permutation test was used to determine whether the experimentally obtained ROC area is significantly greater than what would be expected by chance at p < 0.01 (Britten et al., 1996, see Statistical Analysis).

**Figure 10.**
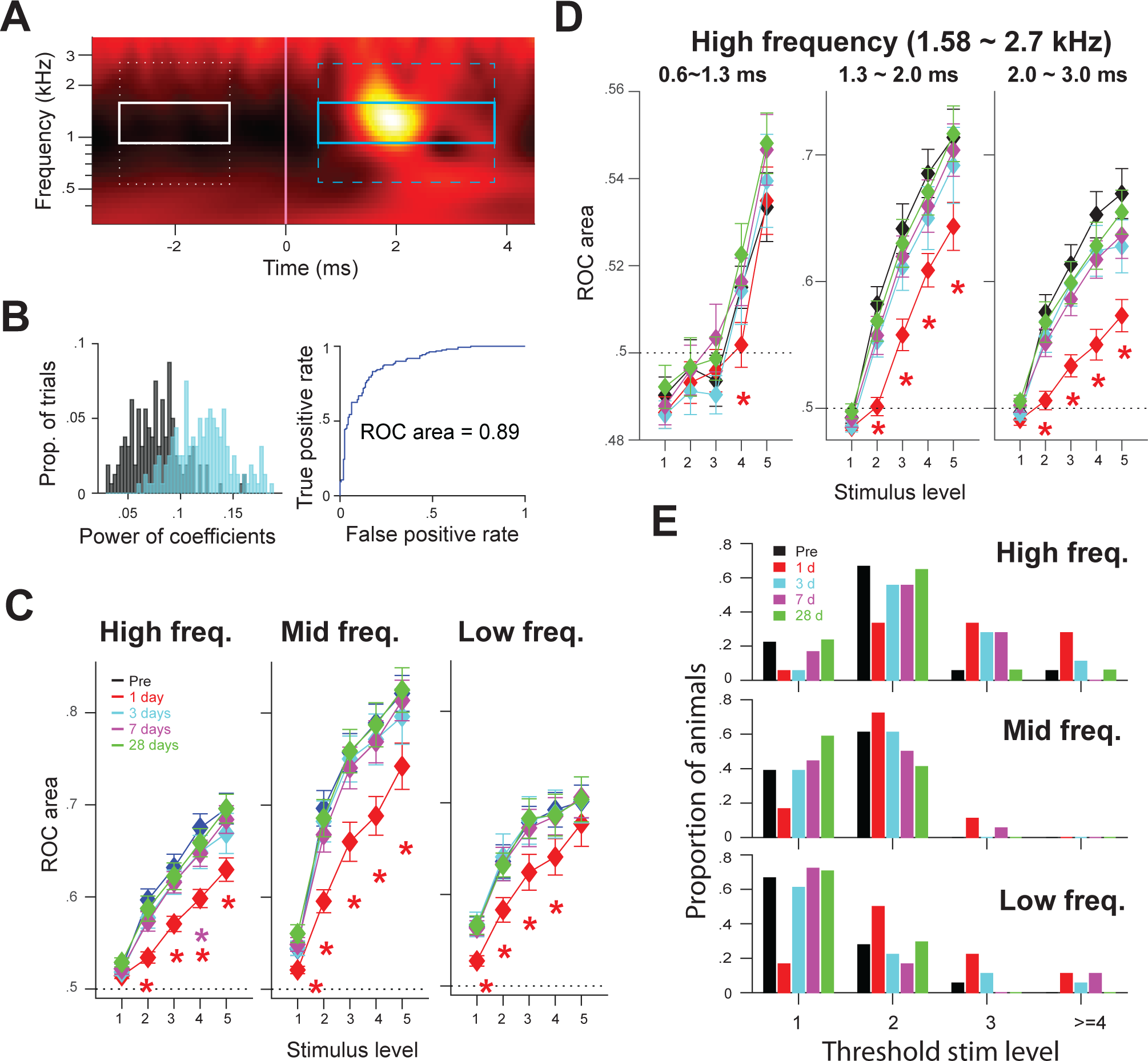
Area under Receiver Operating Characteristics curve (ROC area) was calculated based on the power of single-trial wavelet transform coefficients. **A**: Averaged power of single-trial wavelet transform coefficients in response to 5.5 g/ms jerks is shown as a heat map. Solid, white line indicate the onset of jerks. Boxes prior to the onset of jerks show the frequency x time regions where the “noise” distribution of power is taken for calculation of ROC area at corresponding frequency regions. **B**: Left panel: power of coefficients were averaged across 0.92 ∼1.58 kHz from 0.6∼3.8 ms (indicated by a cyan box in **A**) for each trial, and the distribution across trials is shown in cyan bars. The averaged power across the same frequency range prior to the onset of stimulation (−3 to −1 ms, indicated by a white box in **A**) was also calculated for each trial, and the distribution across trials is shown in black bars. Right panel: Receiver Operating Characteristics (ROC) curve was obtained from the ‘signal’ (cyan bars) and ‘noise’ (black bars) distributions by using decision criteria traversing the power of coefficients and plotting the true positive rate against false positive rate. ROC area is the area under the ROC curve. **C**: The population-average ROC area is plotted as a function of stimulus level based on the power of single-trial wavelet transform coefficients in high (left panel, coefficients averaged in 1.58∼2.7kHz and 0.6∼3.8 ms), mid (middle, 0.92∼1.58 kHz and 0.6 ∼ 3.8 ms), and low frequency regions (right, 0.54∼0.92 kHz and 0.6∼3.8 ms). The corresponding windows are depicted with cyan dashed- and solid-boxes in **A**. Diamond symbols indicate the population mean ROC area, and the error bars indicate standard errors in mean. Asterisk indicates significant difference at post-exposure day compared to pre-exposure with signrank test at p < 0.05. **D**: For the high frequency region, ROC area was calculated in finer temporal windows: 0.6∼1.3 ms (top left panel), 1.3∼2.0 ms (top middle), and 2.0∼3.4 ms (top right). Pre-exposure (black), 1 day (red), 3 days (cyan), 7 days (magenta), and 28 days (green) after noise exposure. **E**: Distribution of minimum stimulus level where ROC area was significantly different from what is expected by chance with a permutation test at p < 0.01. Threshold based on ROC areas in high, middle, and low frequency ranges (using time window of 0.6 ∼ 3.8 ms) are shown on the top, middle, and bottom rows, respectively.

The ROC area based on the power in high-, mid-, and low-frequency WT coefficients area is compared between exposure days (Fig. 10C), and shows a significant reduction in signal detectability on 1 day after exposure and recovery by 28 days (high-frequency: p = 0.0006, 0.0029, 0.0014, and 0.0014 for stimulus levels 2∼5 at day1 by signrank test, p = 0.048 for stimulus level 4 at day 7; mid-frequency: p = 0.0249, 0.0007, 0.0038, 0.0033, and 0.0222 for stimulus levels 1∼5, respectively, at day 1, 0.57 and 0.0065 for stimulus levels 1 and 2 at day 1; low-frequency: p = 0.0033, 0.0198, 0.0156, 0.0279 for stimulus levels 1∼4, respectively, at day 1). For the high-frequency region, where the temporal precision is highest, the ROC area was also calculated in 3 small time windows (Fig.10D). For earliest window (0.6∼1.3 ms), where the activity relates most to the P1-N1 component, the ROC area was much smaller and closer to 0.5 (chance) compared to the other time windows that correspond to the P2-N2 or P3-N3’ components. ROC areas were below 0.5 in many animals at lower stimulus levels (in some animals, significantly below 0.5 with a permutation test), indicating the power of the high-frequency wave during that time period is suppressed compared to the background. This result is consistent with the idea that the P1-N1 component may be better represented by the timing aspect of the high-frequency waves in the VsEP rather than their power.

We determined for each animal a threshold stimulus level, defined as the minimum stimulus level yielding a significant ROC area (by a permutation test at p < 0.01; see Statistical Analysis in Materials and Methods) based on the power of the WT coefficients in high, middle, and low frequency ranges. We found that threshold stimulus levels were smallest for the low frequency region (Fig.10E), implying that the power of the low-frequency wave is more sensitive for detection of a jerk compared to the power of higher frequency waves. Despite this difference in sensitivity, population thresholds determined in all three frequency regions increased 1 day after exposure and then returned to pre-exposure level within 28 days after noise exposure (Fig.10E).

#### VsEP response to positive- and negative-polarity jerks

The positive and negative jerks likely excite different populations of afferents, so we examined SI and normalized power of WT coefficients of the separate waveforms produced by each jerk polarity. Positive- and negative-jerks induce slow, stimulus-locked waves in opposite directions, which cancel out when the waveforms are averaged (Fig.1A). The stimulus-locked wave is characterized by an increase in both SI and normalized power at the onset of stimulus (denoted with white arrow heads on right most column on Fig.11 and 12), that becomes stronger for higher stimulus levels. It appears in all frequency regions but is strongest in the lowest frequency region. Because the temporal precision of a wavelet transform is poorest in the low frequency region (larger temporal durations of wavelets having lower center frequencies), the effect of the stimulus-locked wave smears over a wider time range. To our surprise, SI or normalized power related to the stimulus-locked waves, which has been considered as the electro-magnetic artifact due to the shaker, showed a significant increase 28 days after exposure compared to baseline. This implies that at least some part of this slow wave might have a biological origin modulated by noise exposure.

**Figure 11.**
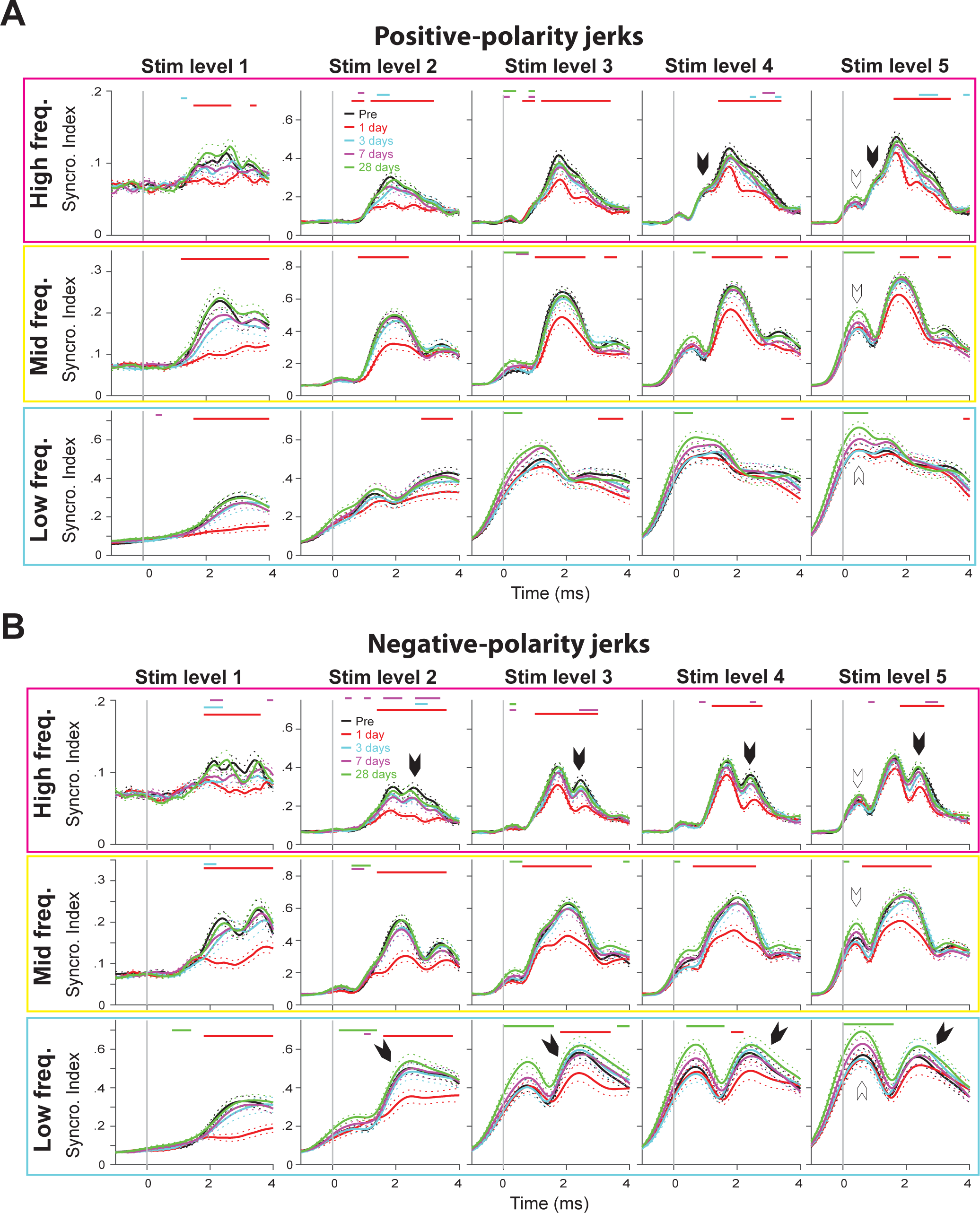
Population-average SI of wavelet transform coefficients for positive-(**A**) and negative-jerk response (**B**). Presented as in Figure 5A.

**Figure 12.**
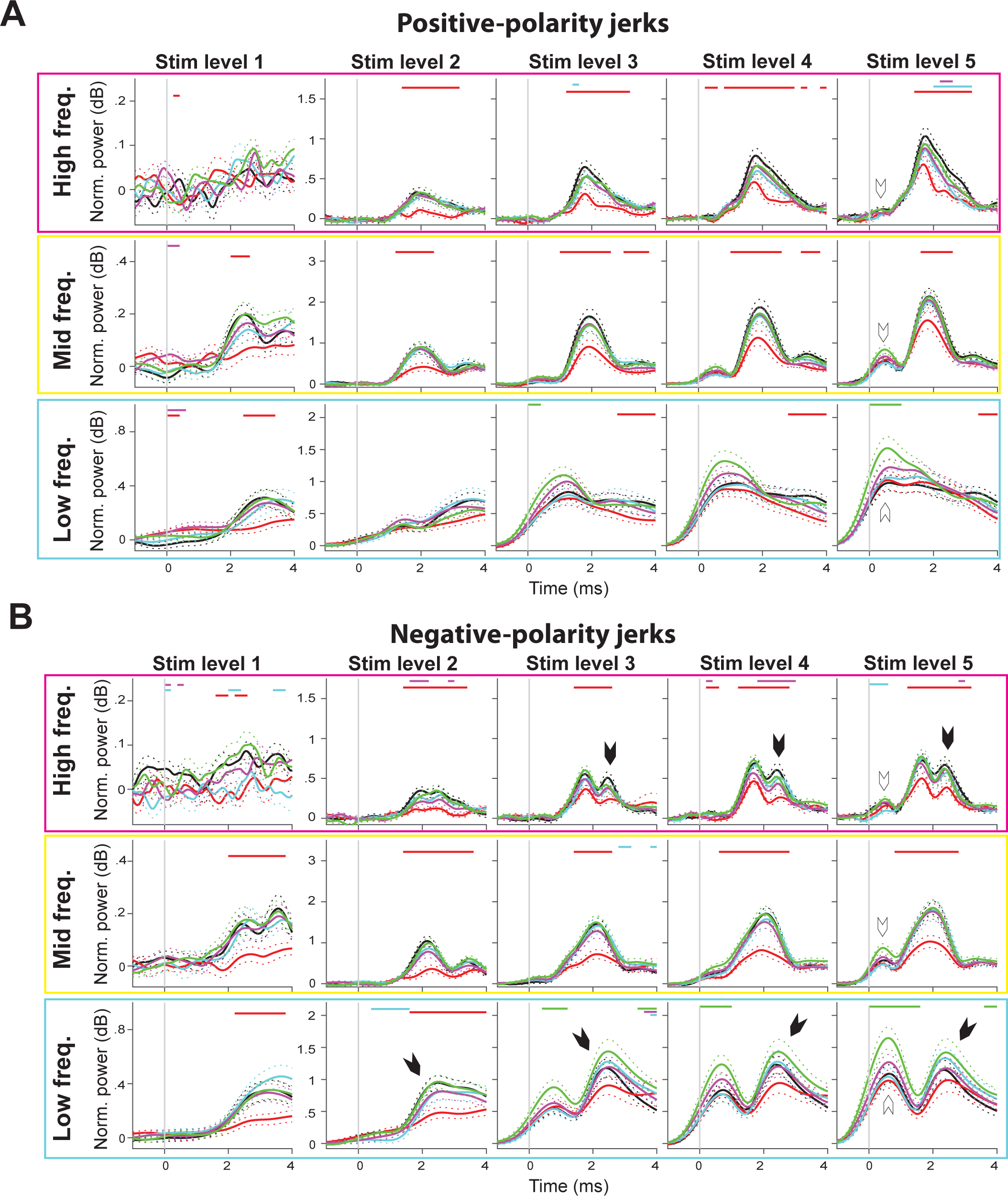
Population-average normalized power of wavelet transform coefficients for positive-(**A**) and negative-jerk response (**B**). Presented as in Figure 9A.

The time-frequency analysis of VsEP waveforms in response to positive- and negative-jerks shows a difference in the pattern of the responses. For example, both SI and normalized power in the high frequency region show a more distinct secondary peak corresponding to P3-N3’ dip for the negative-jerk response compared to the positive jerk (black arrow heads on the top row panels in Fig.11B and 12 B). Similarly, low-frequency SI and normalized power have a more pronounced peak around 2 ms, likely corresponding to a slow P3-N3 component of the VsEP, in the negative-jerk response (black arrow heads on the bottom row panels in Fig.11B and 12 B). The high-frequency SI for the positive-jerk response shows a more pronounced inflection point near the time of P1-N1 peak than for the negative-jerks (black arrow heads, two left most panels on top row of Fig.11A). Even with these differences in temporal pattern, the effect of noise exposure described for average-polarity responses holds for positive- and negative-polarity responses.

### Reduced number of calretinin-positive calyces following noise exposure

Calretinin is a calcium-binding protein expressed in the terminations of calyx-only afferents that innervate striolar/central type-1 hair cells in the vestibular end-organs. It is not expressed in the terminations of dimorphic afferents that innervate both type-1 and type-2 hair cells. Otolith irregular afferents, comprised of calyx-only and striolar dimorphic afferents, are sensitive to high-frequency changes in linear acceleration of the head (Goldberg et al., 1990; Jones et al., 1998; Jones et al., 2011; Jamali et al., 2019). Thus both classes of afferent may respond to jerk stimuli and contribute to the VsEP. Additionally, irregular afferents in the saccule steeply increase their firing rate in response to air conducted sound above 80 dB (Curthoys et al., 2016), and are likely susceptible to noise exposure.

We assessed calretinin expression immunohistologically in saccules harvested 28 days after an exposure to 110 dB noise. We found that the number of calretinin-positive (CR+) calyces in noise-exposed animals was significantly reduced compared to a normative set of animals in a similar age range that were not exposed to noise (p = 1.13−10^-4^ with two-sample t-test, Fig.13A and B). The number of CR+ calyces in the saccule was also significantly correlated with the reduction of SI in high, mid, and low frequency regions in response to the smallest-amplitude, positive-polarity jerks (0.32 g/ms) 1-day after noise exposure (high-frequency: r = 0.68, p = 0.014 by Pearson’s correlation test; mid-frequency: r = 0.63, p = 0.028; low-frequency: r = 0.66, p = 0.021; left most column in Fig 14A). A significant correlation between the number of CR+ calyces and high-frequency SI persisted up to 7 days after exposure before becoming statistically insignificant 28 days after noise exposure (r = 0.74, p = 0.0057 at day3; r = 0.72, p = 0.0083 at day7; r = 0.40, p = 0.20 at day 28; top row in Fig.14A). A possible correlation between the number of CR+ calyces and SI in response to negative-polarity jerks (Fig.14C) appears weaker (only significant for high-frequency on day 3, r = 0.63, p = 0.030) than for response to positive-polarity jerks. In addition, the loss of CR+ calyces in the saccule appears to be tied more strongly to the reduction in higher-frequency SI than in the lower frequency.

**Figure 13.**
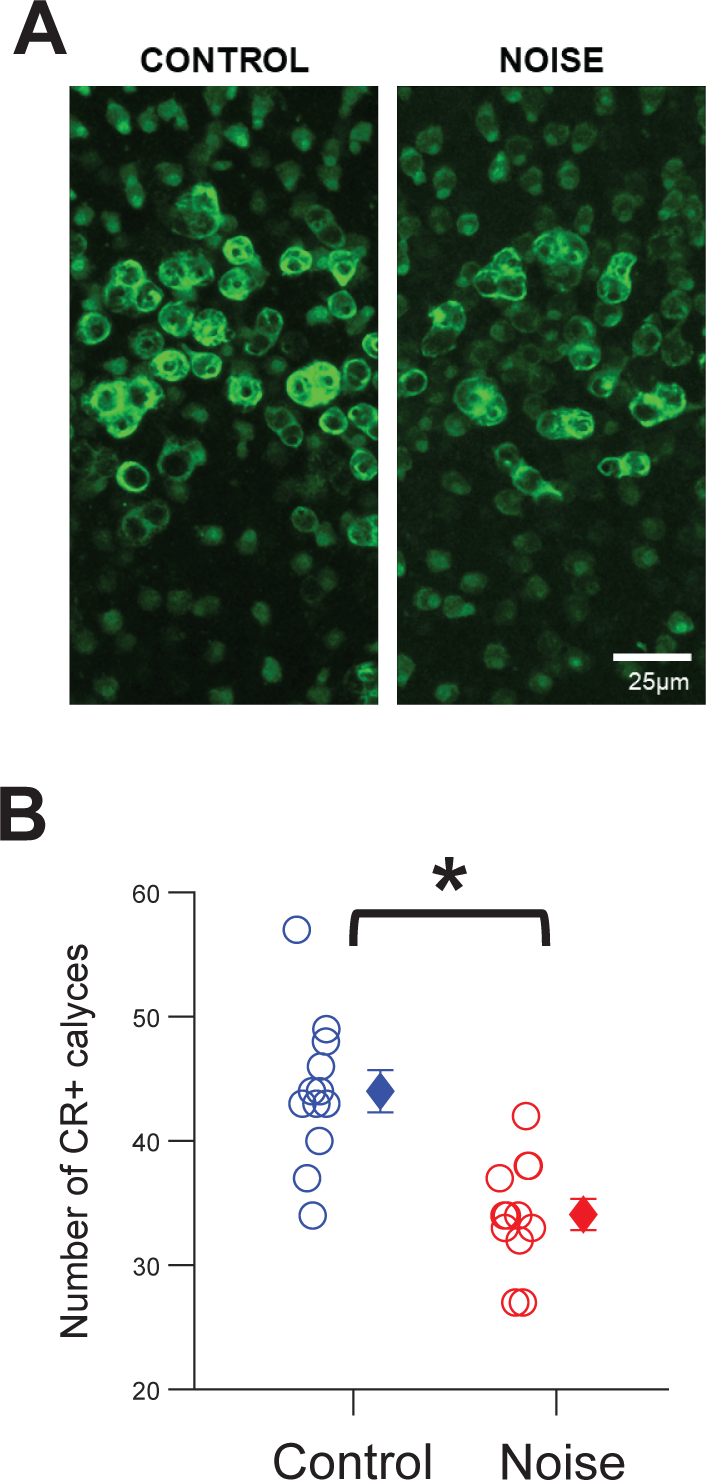
**A:** Two-dimensional z-stack-collapsed images of calretinin immunoreactivity in the saccule of a control (left panel) and a noise-exposed animal (right panel). **B:** Number of CR+ calyces in the saccule of control animals (blue) and noise-treated animals (red). Each circle denotes a count from each animal, a diamond symbol denotes a population mean, and error bars are SEM. Asterisk denotes significant change in mean by t-test (p = 1.13−10^-4^).

**Figure 14.**
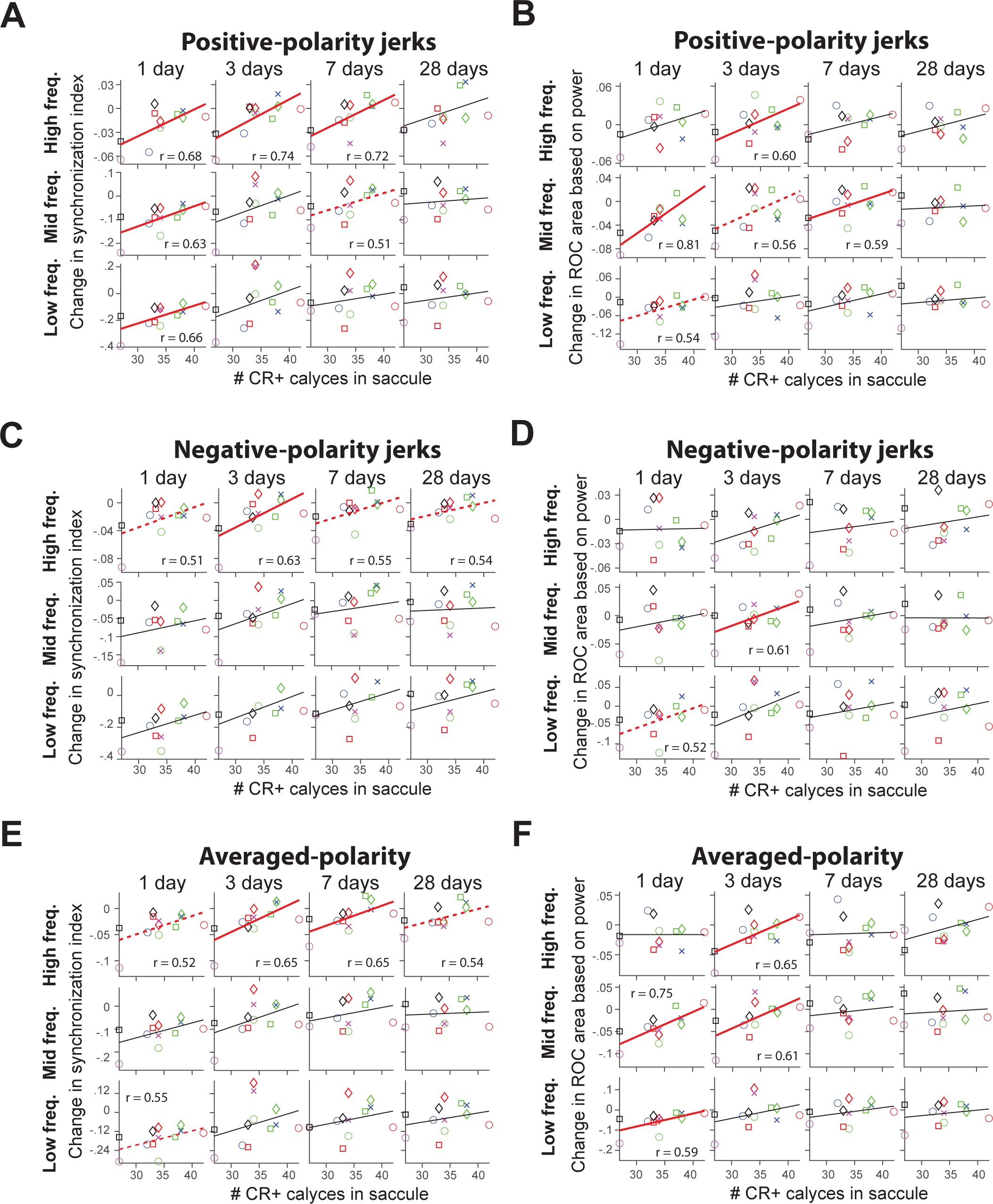
**A**: Scatter plots of the number of calretinin-positive (CR+) calyces in the saccule and the change in SI of wavelet transform coefficients in response to positive jerks at the smallest amplitude (0.32 g/ms). Synchronization-index (SI) was averaged over a time window of 1.0∼3.5 ms for high (1.58∼2.7 kHz, top row panels), 1.0∼3.5 ms for mid (0.92∼1.58 kHz, mid row panels), and 1.8∼3.5 ms for low frequency regions (0.54∼0.92 kHz, bottom row panels). The time window was set so that SI due to the ‘artifact’ will not be included in the analysis. The change in SI was calculated as SI on 1 day (left most column), 3 days (2^nd^ column from the left), 7 days (3^rd^ column from the left), or 28 days after exposure (right most column) minus pre-exposure SI. Linear fits are shown as a red solid line for significant correlations (p < 0.05) with Pearson’s correlation test. Dotted red lines denote marginally significant correlations (p < 0.10). Each combination of symbol and color designates an individual animal’s data point and is consistently used for all other panels. **B**: Scatter plots of the number of CR+ calyces in the saccule and the change in ROC area based on the power of WT coefficients in response to positive jerks at the smallest amplitude (0.32 g/ms). ROC area was calculated by averaging the power of WT coefficients over a time window of 1.0∼3.8 ms for high (1.58∼2.7 kHz, top row panels), 1.0∼3.8 ms for mid (0.92∼1.58 kHz, mid row panels), and 1.8∼3.8 ms for low frequency regions (0.54∼0.92 kHz, bottom row panels). Average power over a time window of −0.3∼−0.1 ms prior to the onset of jerk across the corresponding frequency range in every trial was used as ‘noise’ distribution of power for ROC area calculation. **C** and **D**: Same as **A** and **B**, respectively, but for negative-jerk response. **E**: Same as **A**, but for polarity-averaged response. SI was averaged over a time window of 0.6∼3.8 ms for high, mid, and low frequency regions. **F**: Same as **B**, but for polarity-averaged response. Power of WT coefficients was averaged over a time window of 0.6∼3.8 ms for high, mid, and low frequency regions.

We also found a significant correlation between the number of CR+ calyces and a change in ROC area based on the power of WT coefficients in response to positive-polarity jerks on 1, 3, and 7 days, but not 28 days after noise exposure (Fig.14B; high-frequency: r = 0.60, p = 0.039 at day 3; mid-frequency: r = 0.81, p = 0.0015 at day1, r = 0.59, p = 0.042 at day7). Similar or slightly reduced correlations were seen for negative-polarity jerks (Fig.14D; mid-frequency: r = 0.61, p = 0.037 at day3).

At higher jerk levels 2∼5, neither the change in SI nor power-based ROC areas were significantly correlated with the number of CR+ calyces in the saccule.

## Discussion

We examined the effect of noise exposure on the VsEP using a time-frequency analysis with a wavelet transform on a single-trial basis. Our results reveal that exposure to 110 dB noise for 6 hours significantly reduces an animal’s vestibular sensitivity to jerk 1 day after the exposure. However, the loss of sensitivity is transient and quickly recovered to a pre-exposure level by 28 days after exposure on a population level. The wavelet transform of single-trial VsEP waveforms allowed us to examine the variability in the power/strength of waves underlying single-trial VsEP waveforms as well as the temporal precision of these waves across trials. This is information that cannot be obtained by examination of trial-averaged VsEP waveforms. Our results showed that 110 dB noise exposure temporarily undermines both the power and temporal precision of the waves underlying the VsEP.

We found that the mean power of the single-trial WT coefficient was significantly reduced 1 day after exposure, and the size of the reduction was greater at smaller jerk levels, especially for the low frequency VsEP waves (Fig.9A). The trial-by-trial variability in the power following the onset of a jerk was also significantly greater 1 day after exposure (Fig.9B). As a result, signal detectability of head jerks based on the power of single-trial WT coefficients, as measured by ROC area, was significantly reduced over the same time frame (Fig.10C). Similarly, SI of WT phase, which measures the degree of coherence in timing of waves across trials, was significantly reduced on 1 day after exposure (Fig.5A). The degree of reduction in SI was larger at smaller jerk levels for all frequency regions. Each of these measures, SI and the mean/variability in the power of WT coefficients, recovered to pre-exposure levels on a population level during the 3∼28 days after exposure.

### Afferents with calretinin-positive (CR+) calyces and VsEP

Our previous reports showed that exposure to 120 dB noise for 4∼6 hours significantly reduced the amplitude of the VsEP waveform 28 days after exposure as well as the number of CR+ calyces in the saccules (Stewart et al., 2020) and utricles (Niwa et al., bioRxiv). Reduction in the number of CR+ calyces was significantly correlated with the degree of reduction in VsEP amplitude (Niwa et al., bioRxiv). Those results appeared to support a functional role of afferents with CR+ calyces in generation of the VsEP, which was consistent with their known high sensitivity to linear forces acting on the head (Goldberg, 2000 for review) as well as their likely susceptibility to air conducted sound above 100 dB (Curthoys et al., 2016).

In the present study, we found that the number of CR+ calyces counted in tissues collected 28 days after exposure to 110 dB noise was also significantly reduced compared to control, indicating that a morphological abnormality persists whilst physiological functionality, as measured by the VsEP, returns to baseline. Intriguingly, the number of CR+ calyces in the saccule was significantly correlated with a reduction in SI or signal detectability (ROC area) of the smallest-amplitude jerk, which was most affected by noise exposure. The correlation was significant on earlier days (1, 3, or 7), but not 28 days after exposure. This result suggests the morphological variability among animals 28 days after noise exposure is not significantly correlated with the individual variability in VsEP measured on 28 days after exposure (the same day as tissue collection), but rather related to effects occurring in the first week after exposure.

Our present finding is at odds with a simple interpretation that the decrease in the number of afferents with CR+ calyces leads to a reduced VsEP. Although exposure to 110 dB vs 120 dB noise causes temporary vs permanent changes in the VsEP respectively, the number of CR+ calyces was significantly reduced for either exposure level in similar proportions (20∼25% reduction). One interpretation is the number of CR+ calyces (assessed 28 days after both types of exposures) serves as a persistent proxy for the magnitude of noise-induced damages following either 110 or 120 dB nose exposure; however, the reduction in calretinin-positive calyces is not itself directly related to the attenuation of the VsEP.

Our result raises questions about the functional role of calretinin in the terminations of calyx-only afferents. In future studies, the expression level of calretinin in remaining CR+ calyces, as well as the size and complexity of surviving calyx-only terminals and striolar calyces arising from dimorphic afferents, in addition to the number of CR+ calyces, will need to be quantified following noise exposure to explain the differential effect of noise exposure levels. Noise-induced changes in the expression of calbindin, another calcium-binding protein found in calyx-only and dimorph afferents is also possible and should be examined in future experiments to evaluate irregular dimorphic afferents’ susceptibility to 110 vs 120 dB noise exposure and their possible contribution to VsEP waveforms.

The present study shows that the vestibular periphery sustains a morphological alteration for at least 28 days after noise exposure, which is not reflected by the VsEP on a population level. We hypothesize that the vestibular periphery is vulnerable to a single exposure of 110 dB noise, despite the observation of a normal VsEP 7 days after exposure, and that the VsEP might not recover to baseline upon repeated noise exposure. A future study will investigate whether a repeated exposure to 110 dB noise with a 7-day intervals results in a permanent reduction in VsEP as observed after a single 120 dB exposure (Stewart et al., 2020).

### VsEP in response to positive- and negative-polarity jerks

A time-frequency analysis of the VsEP waveform in response to positive- and negative-jerks revealed a difference in their temporal pattern. For example, negative-jerk responses showed a more distinct secondary peak corresponding to P3-N3’ dip than did positive-jerk responses, while positive-jerk responses showed a more pronounced inflection point around the time of the P1-N1 peak than did negative-jerk responses. These differences are not surprising since positive- and negative-jerks likely stimulate different sets of hair cells on either side of the striola in the otolith organs and/or different contributions of utricular vs saccular hair cells. Indeed, a correlation between the number of CR+ calyces in the saccule and reduction in SI or ROC area appeared qualitatively stronger for positive-jerk responses compared to negative-jerk responses. The wavelet transform allows us to separate the underlying waves of the VsEP by frequency so that we can discern the effect of ‘artifact’ over the higher frequency regions without resorting to polarity-averaging. This approach may be useful to determine which population of otolith neurons contribute to positive and negative jerks in VsEP stimulation.

Positive- and negative-jerk responses on a single-trial basis may also be useful for understanding the role of different components of the VsEP in terms of serial/parallel processing in the vestibular system. The wavelet transform of single-trial VsEP waveforms allows us to determine the trial-by-trial correlation between strength and well-timed activity (how well a phase of a given trial aligns with the preferred phase across all trials) at different time and frequency points during the response. For example, does stronger and/or more precisely-timed P1-N1 activity lead to larger P2-N2, P3-N3’, or P3-N3 waveforms in a single trial? Independent analyses of positive- and negative-jerk responses is more useful than polarity-averaged responses because polarity averaging of pairs of positive- and negative-jerk responses (not strictly a single-trial analysis) can obscure possible trial-by-trial correlations.

The P1-N1 component is thought to originate from vestibular primary afferents while P2-N2 and subsequent waves may include secondary vestibular neurons in the brain stem (Nazareth and Jones, 1998). Recently, Pastras et al. (2023) showed that the P1-N1 component is likely a compound action potential comprised of irregular afferents driven by non-quantal transmission from type-1 hair cells to post-synaptic calyceal terminals. This finding raises the question of a possible contribution of action potentials originating from quantal-transmission to P1N1 or later peaks of the VsEP. It is also possible that action potentials from quantal-transmission occur only in those trials where non-quantal transmission fails to evoke an action potential. Analysis of the VsEP on a single-trial basis could provide additional insights into what the different components of the VsEP represent, and the wavelet transform is one way of quantifying noisy single-trial waveforms.

## Acknowledgements

The study was supported by NIH: R01DC018003, R01AG073157, VA: I01RX003250, IK2RX003271.

